# DNA methylation correlates with lifespan and predicts remaining lifespan in the zebra finch

**DOI:** 10.64898/2026.09.10.750555

**Authors:** Marianthi Tangili, Per J. Palsbøll, Simon Verhulst

**Affiliations:** Groningen Institute for Evolutionary Life Sciences, University of Groningen, Groningen, The Netherlands; Center for Coastal Studies, Provincetown, Massachusetts, USA

**Keywords:** epigenetic clock, aging, zebra finch, lifespan variation, lifespan prediction, developmental conditions, longitudinal

## Abstract

Biomarkers that correlate with age are essential tools for understanding aging and lifespan variation. DNA methylation (DNAm) changes predictably with age in parts of the genome. In humans, epigenetically “old” individuals relative to their chronological age also have a reduced life expectancy but whether the link between epigenetic age and lifespan is a general feature remains an open question. We explored age-related changes in DNAm in the zebra finch (*Taeniopygia castanotis*), a key avian model species, using 100 longitudinal whole genome methylomes from 50 captive adults monitored until their natural death. We found genome-wide hypomethylation with age, with DNAm decreasing faster in individuals with shorter lifespans and identified 29 CpG sites where DNAm changed significantly with age. We developed an epigenetic aging clock based on 119 CpG sites, that predicted chronological age with high accuracy (median absolute deviation=0.68 years, 8.2% of maximum lifespan in our dataset). Females raised in large broods, which have shorter lifespans, showed increased epigenetic age acceleration, consistent with faster biological aging. However, epigenetic age acceleration did not predict lifespan or remaining lifespan. In contrast, the within-individual rate of change in epigenetic age significantly predicted lifespan: faster epigenetic aging was associated with a shorter lifespan. Moreover, a “doom” clock, trained to predict post-sampling lifespan, successfully predicted remaining lifespan. Our findings provide the first evidence that DNAm provides a window into biological aging in birds, showing how early-life environments shape the aging trajectory via the epigenome and underscore the value of longitudinal data in aging studies.

## Introduction

Aging rates vary substantially between individuals, populations and taxa (Lemaître et al., 2024; Lowsky et al., 2014), with individuals of the same chronological age exhibiting large variation in morbidity and mortality. Consequently, there is growing interest in quantifying an individual’s biological age, a measure aiming to capture an individual’s somatic integrity, as a means towards uncovering the mechanisms of aging that drive variation in aging rates and lifespan, potentially improving clinical practice (Lee, 2026). The development of aging clocks, machine learning models trained on molecular features (e.g. metabolomics, proteomics, transcriptomics, microbiomics and epigenomics) to estimate age has revolutionized our understanding of the molecular underpinnings of aging (Rutledge et al., 2022).

DNA methylation (DNAm), an epigenetic modification that involves the addition of a methyl group to a cytosine catalyzed by methyltransferases, defines and maintains cell identity (Bogdanović & Lister, 2017) through regulation of gene expression (Moore et al., 2013). Genome-wide levels of DNAm decrease with age in vertebrates (Unnikrishnan et al., 2018); a hypomethylation that can lead to tumorigenesis (Berman et al., 2015; Timp et al., 2014) and genomic instability (Karpf & Matsui, 2005; Sheaffer et al., 2016) which along with deregulation of gene expression are known hallmarks of aging (López-Otín et al., 2023).

Epigenetic clocks leverage age-related changes in DNAm on specific CpG sites across the genome to predict chronological age with high accuracy (Horvath, 2013). In humans, individuals with an elevated epigenetic age compared to their chronological age have a higher mortality risk and consequently a shorter lifespan (Kuo et al., 2026; Levine et al., 2018; Lu et al., 2019). Thus, epigenetic age captures meaningful variation in somatic integrity and lifespan beyond chronological age, positioning it as a potential tool for clinical and conservation applications.

Epigenetic clocks of age have been developed in a variety of non-model species(Bors et al., 2021; Tangili et al., 2023; Thompson et al., 2017), including birds (Wolf & Tangili, 2026), and are increasingly used in ecology and evolution (Parrott & Bertucci, 2019; Tangili et al., 2023). However, the extent to which epigenetic aging indicates lifespan variation in non-human species remains unresolved. Validation of epigenetic clocks as a predictor of lifespan beyond humans would increase the potential of epigenetic clocks as a tool in experiments on wild and captive animals to investigate how environmental, social and life-history factors shape individual aging trajectories. Indeed, a handful of studies illustrated the potential of this approach. For example, in baboons (*Papio cynocephalus*), a high social rank was associated with accelerated epigenetic aging, but not mortality (Anderson et al., 2021). Male bottlenose dolphins (*Tursiops aduncus*) with stronger social bonds had lower epigenetic age estimates (Gerber, Peters, et al., 2025). In birds, nestling great tits (*Parus major*) raised in experimentally reduced broods were epigenetically older compared to control individuals, reflecting faster development in reduced broods (Haller et al., 2025), and, beyond age prediction, DNAm patterns can predict other traits of interest such retroactively quantifying aspects of early-life conditions (Tangili, Palsbøll, et al., 2026). However, whether epigenetic age acceleration predicted mortality or life expectancy in these systems remains an open question.

The main goal of this study was to characterize age-related changes in DNAm to develop an epigenetic clock of age in the zebra finch (*Taeniopygia castanotis*). We tested if epigenetic aging predicted lifespan variation in this non-mammalian vertebrate. Extending epigenetic aging research to avian species holds great promise for comparative studies aimed to identify aging processes shared across vertebrates among a diverse array of life-history strategies. In addition, similar sized birds are long-lived compared to mammals (Harper & Holmes, 2021). Moreover, insofar that clocks are based on blood samples, using birds has the advantage of using a single cell type, erythrocytes, which are nucleated in birds, as in amphibians and reptiles. In contrast, mammal erythrocytes are not nucleated and hence DNA is obtained from leukocytes, and variation in cell type composition potentially biases age estimation (Aviv & Verhulst, 2025).

The genome-wide hypomethylation hypothesis of aging posits that an overall decrease in DNAm with age relaxes the epigenetic control of gene expression thus driving aberrant transcription that contributes to aging (Unnikrishnan et al., 2018). To investigate this hypothesis, we quantified genome-wide levels of DNAm, and how it changed with age. We then developed an epigenetic clock of age. We complemented this analysis with identification of CpG sites whose DNAm significantly changes with age, which allowed us to subsequently assess how DNAm at these CpG sites contributed to the epigenetic clock of age.

Our study was based on a longitudinal whole-genome DNAm dataset from 50 adult known-age zebra finches of both sexes that were part of a long-term experiment, where individuals’ developmental history was manipulated by being reared in either small or large broods. The experiment revealed that being female, or having been reared in a large brood, both (independently) resulted in a reduced lifespan (Briga et al., 2017). To investigate the association between the epigenetic clock estimates and the phenotype, we compared epigenetic age acceleration between sexes and brood sizes. Furthermore, to test the hypothesis that epigenetic aging drives the observed differences in lifespan observed in our study population, we compared epigenetic age acceleration and within-individual changes in epigenetic age with lifespan. Lastly, we developed an epigenetic clock of remaining lifespan, as this may yield a more direct link between the methylome and remaining life expectancy compared to the age acceleration metric.

## Methods

### Subjects and brood size manipulation

The present data set consists of birds from the experiment reported in (Briga et al., 2017), where background and treatment of subjects is described in detail. In brief, adult zebra finches were randomly matched and provided with nest boxes and nesting material. Nestboxes were checked daily for the presence of eggs or chicks. Chicks were cross-fostered at age five days or before to broods of either two or six chicks, which is the range of brood sizes observed naturally in the wild (Zann, 1996) and in captivity (Griffith et al., 2017). Siblings were assigned to different broods, and parents never reared their own offspring. After reaching adulthood (≥100 days), birds were moved to outdoor aviaries, each containing single sex groups of 18–24 adults where they were followed for the rest of their natural lifespan.

DNAm was measured in blood samples collected from the brachial vein and stored in -80°C in glycerol buffer from 50 individuals distributed equally over the small and large broods (25 raised in small, 25 raised in large broods) and almost equally over the two sexes (26 males, 24 females). In the study by Briga et al. (2017), foraging costs in the outdoor aviaries were either benign or harsh (see Koetsier & Verhulst, 2011), but the present data set consists of birds in the harsh foraging conditions only, because only in these conditions did rearing brood size impact lifespan (Briga et al., 2017). DNA methylation data were extracted for two samples per individual collected early and later in their life (Fig.S1a), with an average sampling interval of 995 (S.D.=742) days. For more information on sample information and experimental design, see Tangili et al. (2026).

### Enzymatic Methyl-seq data

We extracted DNA according to the manufacturer’s protocol using innuPREP™ DNA Mini Kit (Analytik Jena GmBH) from 3 uL of erythrocytes. Next-generation sequencing was conducted by the Hospital for Sick Children (Toronto, Canada). The amount of DNA was quantified using a Qubit High Sensitivity Assay (Thermo Fisher Scientific). 200 ng of DNA was used as input material for library preparation using the Next Enzymatic Methyl-seq™ Kit (New England Biolabs Inc.) following the manufacturer’s instructions. Briefly, DNA was fragmented by sonication to an average length of ∼500 base pairs (bp) using a Covaris LE220 (Covaris Inc.). Fragmented DNA was end-repaired and ligated to Illumina sequencing adapters. The 5-methylcytosines and 5-hydroxymethylcytosines were oxidized by TET2 and cytosines were deaminated by APOBEC. Methyl-seq libraries were subjected to five PCR amplification cycles. Libraries assessed on a Bioanalyzer™ DNA High Sensitivity chip (Agilent Inc.). The amount of DNA was quantified by quantitative PCR using the Kapa Library Quantification Illumina/ABI Prism™ kit according to the manufacturer’s instructions (La Roche Ltd.). Libraries were pooled in equimolar amounts and sequenced to a minimum of ∼100M paired-end reads (2 × 150bp) per sample on a 10B flow cell using a NovaSeqX™ (Illumina Inc.) platform.

Sequences were trimmed using Trim Galore! v0.6.10 (Krueger et al., 2023) in paired-end mode. The data were assessed before and after trimming using FastQC v. 011.9 (Andrews, 2010) and MultiQC v. 11.14 (Ewels et al., 2016). Trimmed reads were aligned against the *in silico* bisulfite converted zebra finch reference genome (GCA_003957565.4, (Rhie et al., 2021) using Bismark v. 0.14.433 (Krueger & Andrews, 2011) using the Bowtie 2 v. 2.4.5 alignment algorithm (Langmead & Salzberg, 2012). The average mapping efficiency was at 65.14% (SD: 2.79). Mean autosomal DNAm did not appear to be affected by the storage time (range 8-16 years) of the samples (estimate= 0.03 ± 0.04, t = 0.85, p = 0.39).

### CpG site filtering

After DNAm extraction we ended up with 19,995,101 unique CpG sites. Using the R package *Methylkit* v.1.18.0 (Akalin et al., 2012), we removed CpG sites with a coverage of less than 5 (Ziller et al., 2015) and maximum coverage at the 99.9th percentile of coverage, which resulted in 15,797,606 sites. Moreover, we removed CpG sites that were not shared between all samples, yielding 1,120,977 sites. Finally, CpG sites with DNAm 0 or 100% in all samples were removed, resulting in a final data set of 1,141,144 CpG sites available to train the epigenetic clock on. In the remainder of the manuscript we refer to this as “the filtered CpGs”.

### Principal component analysis (PCA)

To access genome-wide patterns of DNAm variation and identify potential batch effects and outliers, we performed principal component analysis (PCA) on the filtered, CpG-level DNAm of the filtered CpGs. PCA was performed using the *prcomp*() function in R with centering and scaling to standardize CpG sites across the dataset.

### Epigenome wide association study (EWAS) of chronological age

We performed an epigenome wide association study (EWAS) of chronological age using the R package WGNA v.1.73 (Langfelder & Horvath, 2008). We utilized the “standardScreeningNumericTrait” function which computes a Pearson’s correlation of the DNAm of the filtered CpGs with chronological age and outputs a z-statistic and corresponding p-value. A false discovery rate (FDR)–adjusted threshold (Benjamini–Hochberg method, Benjamini & Hochberg, 1995) was used to correct for multiple testing.

### Epigenetic clock

To develop the epigenetic clock of age, we used elastic net regression implemented in the *glmnet* v. 4.1-8 package (Friedman et al., 2010). The DNAm values of the 1,141,144 CpG sites were used as predictors while chronological age at sampling was the response variable (with or without log transformation, see below).

The mixing parameter α was optimized by testing values from 0 to 1 in increments of 0.1, performing 100 iterations of 10-fold-cross-validation for each value. As the optimal α we selected the value that yielded the lowest average cross-validated mean squared error. Using the best-performing *α*, we then determined the optimal regularization parameter λ that corresponded to the lowest mean cross-validated error. For the epigenetic clock trained on age the optimal α was 1 and the optimal λ was 9.74, while for the epigenetic clock trained on the logarithmic transformation of age they were 0.6 and 0.16 respectively.

To evaluate model accuracy while accounting for multiple samples per individual, we used a leave-one-individual-out cross-validation (LOIOCV) approach. In each iteration, the two samples collected at different ages from one individual were excluded from the model. Model performance was quantified by the Pearson’s correlation of chronological to epigenetic age and the median absolute deviation (MAD) between chronological and epigenetic age. We here consider MAD as the more effective measure of accuracy (Tangili et al., 2023). After validation, a final elastic net model was trained using all samples with the selected *α* and *λ*.

### Statistical analyses

To calculate within-individual mean autosomal DNAm repeatability, also known as intra-class correlation coefficient (ICC), we used the package rptR v.0.9.22 (Stoffel et al., 2017) with mean autosomal DNAm as the response variable, delta age (the difference between an individual’s chronological age at the time of sampling and its average age calculated for both samplings, as fixed effect and individual as a random effect with 1,000 bootstrapping iterations.

To explore whether the genomic distribution of CpG sites increasing vs. decreasing in DNAm with age differed, a contingency table of correlation direction by genomic category was constructed, and a chi-square test of independence was used to evaluate whether the distributions differed between positively and negatively corelated CpGs. To identify which genomic categories contributed most strongly, standardized Pearson residuals from the choi-square test were calculated, with positive residuals indicating enrichment and negative residuals indicating depletion relative to the expected frequencies.

To access whether clock CpGs were distributed equally across genomic categories, enrichment or depletion of clock CpGs relative to the background set of all filtered CpGs was evaluated using Fisher’s exact tests performed separately for each genomic category. In addition, observed vs. expected numbers of CpG sites in each genomic category were compared using binomial tests.

Age acceleration is usually calculated as the residuals of the linear model of epigenetic and chronological age (Horvath et al., 2014). However, because we here have longitudinal data we calculated age acceleration as the residuals of the linear mixed effects model of chronological and epigenetic age with BirdID as a random effect using the package *lme4* v. 1.1.35.1 (Bates et al., 2015). Linear mixed models were then fitted to assess whether epigenetic age acceleration differed significantly between the sexes and between individuals raised in small and large broods.

All statistical analyses were conducted in R v. 4.3.2 (R Core Team, 2023).

## Results

### Genome-wide DNA methylation

We observed significant autosomal hypomethylation with age (Fig.1a). There was no sex effect on either the average autosomal DNAm, or on the rate of change with age (Table 1a, Fig.1a). Mean autosomal DNAm exhibited high within-individual repeatability (Fig.1a, R = 0.79, 95% CI = 0.67-0.88, *p* < 0.001) indicating strong individual-specific stability in autosomal DNAm across samples taken on average more than two years apart. Notably, hypomethylation was modulated by lifespan (Table 1b, Fig.1b), with hypomethylation increasing faster with age in short lived individuals.

**Figure 1.**
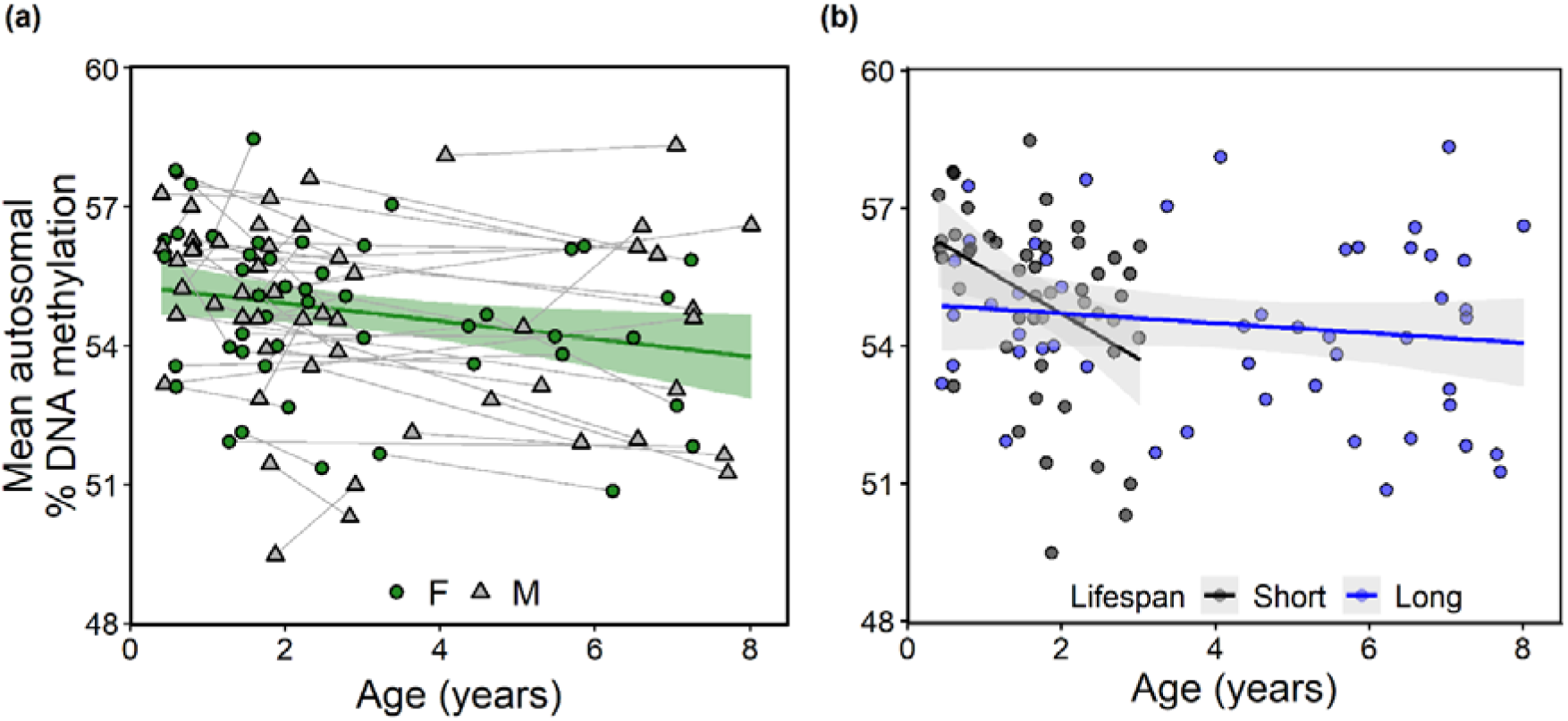
Mean autosomal % DNA methylation in relation to age and lifespan. (a) Relationship of chronological age and mean autosomal DNA methylation. Each point represents one sample of one individual, with lines connecting samples of the same individual. Circles represent females and triangles males. Solid green line represents the fitted linear regression and the shaded area is the 95% CI. (b) Relationship of chronological age and mean autosomal DNA methylation for short- and long-lived individuals. The solid lines represent the fitted linear regression and the shaded areas are the 95% CI. Individuals with “short” lifespans had an average lifespan of 2.44 ± 0.57 years and individuals with “long” lifespans had an average lifespan of 7.04 ± 0.65 years. Individuals were grouped with respect to lifespan for graphical purposes only – in the statistical analysis, summarized in Table 1b, lifespan was entered as continuous variable.

**Table 1.** (a) Mean autosomal % DNA methylation in relation to age and sex. Sex was mean-centered with females coded as -0.5 and males as +0.5 and age is expressed in years. (b) Mean autosomal % DNA methylation in relation to age and lifespan. Age and lifespan are expressed in years.

| **(a)** |  |  |  |  |
| --- | --- | --- | --- | --- |
|  | Mean autosomal DNA methylation |  |  |  |
|  | Estimate (± SE) | t-value | df | Pr(>\|t\|) |
| *Fixed effects* |  |  |  |  |
| Intercept | **55.18 (± 0.30)** | **185.79** | **74.99** | **<0.001 \*\*\*** |
| age (years) | **-0.15 (± 0.05)** | **-3.03** | **56.82** | **0.004 \*\*** |
| sex | -0.27(± 0.59) | -0.45 | 75 | 0.65 |
| age * sex | 0.08 (±0.1) | 0.75 | 56.82 | 0.46 |
| *Random effects* |  |  |  |  |
| bird id | 2.88 |  |  |  |
| Residual | 0.78 |  |  |  |
| **(b)** |  |  |  |  |
|  | Mean autosomal DNA methylation |  |  |  |
|  | Estimate (± SE) | t-value | df | Pr(>\|t\|) |
| *Fixed effects* |  |  |  |  |
| Intercept | **56.46 (± 0.72)** | **78.88** | **87.32** | **<0.001 \*\*\*** |
| age (years) | **-0.68 (± 0.27)** | **-2.53** | **59.87** | **0.01 \*** |
| lifespan | -0.22(± 0.13) | -1.75 | 81.95 | 0.08 |
| age * lifespan | **0.08 (±0.04)** | **2.05** | **60.01** | **0.04\*** |
| *Random effects* |  |  |  |  |
| bird id | 2.79 |  |  |  |
| Residual | 0.74 |  |  |  |

Summarizing DNAm landscapes over all individual CpG sites through principal component analysis did not yield additional insights. The first principal component (PC1) explained 3.5% of the variation in DNAm across samples (Fig.S1b) and showed a near perfect correlation with mean autosomal DNAm per sample (r = 0.97, *p* < 0.001; Fig.S1c), indicating that PC1 captured levels of autosomal DNAm only. The second principal component (PC2) captured 2.2% of the variation in DNAm (Fig.S1b) and was not correlated with age, sex or lifespan (all *p* > 0.1).

### CpG-specific age-related changes in DNA methylation

Pearson’s correlation coefficients between chronological age and DNAm ranged from -0.57 to 0.57 over all CpG sites that passed filtering (N=1,141,144; Fig.2a). Of these CpGs, 62% exhibited negative, and 38% exhibited positive correlations between DNAm and age (Fig.2a). The mean Pearson’s correlation was significantly negative at -0.031 (95% CI = -0.0313, -0.0309). The Pearson’s correlation coefficients within individuals, i.e. between delta age (= chronological age - average age per individual) and DNAm, per CpG site ranged from -0.47 to 0.5 (Fig.S2) and was overall estimated at -0.016 (95% CI = -0.017, 0.016), including zero in the 95% confidence interval. We attribute the difference between the cross-sectional and longitudinal result to the smaller variance in age in the latter analysis. The genomic distribution of CpG sites differed markedly between sites with positive vs. negative correlations of DNAm with age (chi-square test, *p* <0.001). Sites showing increasing DNAm with age were strongly enriched in introns (standardized residual = 23.99) and moderately enriched in promoters (3.12), while sites showing decreased DNAm with age were enriched in intergenic regions (21.52) and exons (6.38).

**Figure 2.**
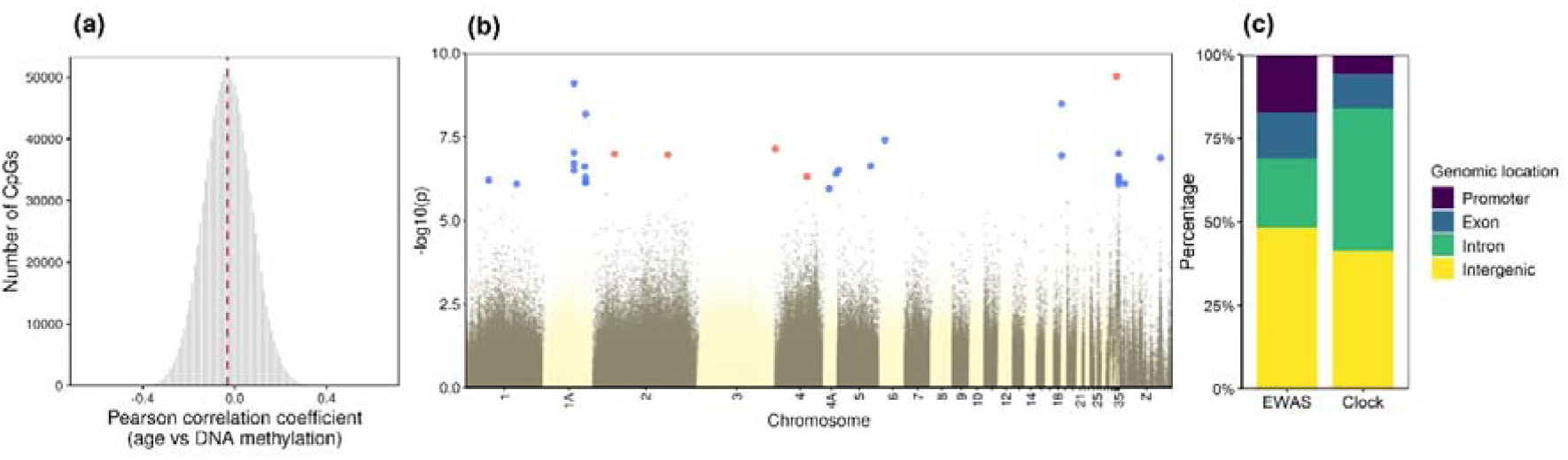
CpG level correlations between methylation and age. (a) Distribution of Pearson correlation coefficients for all CpG sites that passed filtering (including the Z chromosome, N=1,141,144). The dashed vertical line denotes the mean (-0.031, 95% CI = -0.0313 - -0.0309). (b) Manhattan plot of the epigenome wide association study (EWAS) of chronological age in zebra finches. Red dots represent CpG sites whose methylation significantly increased with age (N=5) and blue dots represent CpG sites whose methylation significantly decreased with age (N=24) after correcting for multiple testing (FDR correction). (c) Distribution of significant CpGs from the EWAS of chronological age (N=29) and clock CpGs (N=119) across genomic location categories. There are 18 shared CpG sites between the two sets of CpGs sites.

The EWAS of chronological age identified twenty-nine CpG sites that exhibited significant correlations after FDR correction (Fig.2b, Table S1). The 29 CpG sites were located in 12 unique chromosomes, including the Z chromosome. Among the 29 CpG sites, DNAm increased with age at five CpG sites, and decreased with age at the remaining 24 CpG sites (Fig.2b). EWAS CpGs were for the most part located in intergenic regions (N=14), followed by introns (N=5), promoters (N=5) and exons (N=4; Fig.3). Using the distribution of filtered CpGs across annotation categories to calculate expected counts, binomial tests indicated that promoters were significantly enriched (*p* = 0.001) and introns significantly depleted (*p* = 0.003) for EWAS significant CpGs. The significant EWAS CpGs were located in or near 20 unique genes, ten of which were in uncharacterized LOC genes. Since this analysis is correlational, the relationship between DNAm and chronological age in the identified CpG sites needs to be validated in an independent dataset in order to make conclusions about the causal relationship with the aging process.

### Epigenetic clock of age

We developed an epigenetic clock using chronological age as the response variable, which predicted age with a median absolute deviation (MAD) of 1.08 years while the correlation between chronological and epigenetic age was r = 0.81 (95% CI = 0.73, 0.87, df = 98, p < 0.001; Fig.S2). However, the relationship between chronological and epigenetic age was non-linear, with faster apparent epigenetic aging early in life and slower changes later on (Fig.S3), suggesting transformations of age may yield better predictions. Indeed, using (natural) logarithmic age as the response variable in the training largely removed the non-linear pattern and yielded a more accurate epigenetic clock with a MAD of 0.68 years (r = 0.81, 95% CI = 0.78-0.9, df = 98, p < 0.001; Fig.4a). We next fitted epigenetic age to known age taking individual identity into account as random effect in a mixed model. This confirmed better performance when the epigenetic clock was trained using natural log-transformed ages (AIC difference: -3.52). Individual identity explained a significant proportion of the total variance in both models with the ICC at 13.5% and 27.7% for linear and log-transformed age, respectively. We see the higher ICC when the clock was trained on log-transformed age as confirmation that log-transformed age yields a better fitting clock, and continued further analyses with this clock only. The ICC values highlight that individuals with a high epigenetic age relative to their chronological age early in life also have a relatively high epigenetic age when resampled years later in life. Epigenetic age increased with chronological age in 88% of all individuals (Fig.4b).

The clock contained 119 CpG sites in 27 autosomal chromosomes (Fig. S4a, Table S1). Chromosome 1A included the highest number of clock CpGs (n=24) and showed significant enrichment for clock CpGs (Fig. S4b; 24 observed vs 9.2 expected; odds ratio=3.18, *p*-two-sided < 0.001). Chromosome 7 was significantly depleted in clock CpGs (Fig. S4b; 0 observed vs. 4.6 expected; odds ratio = 0.00, *p*-two-sided = 0.004).

**Figure 4.**
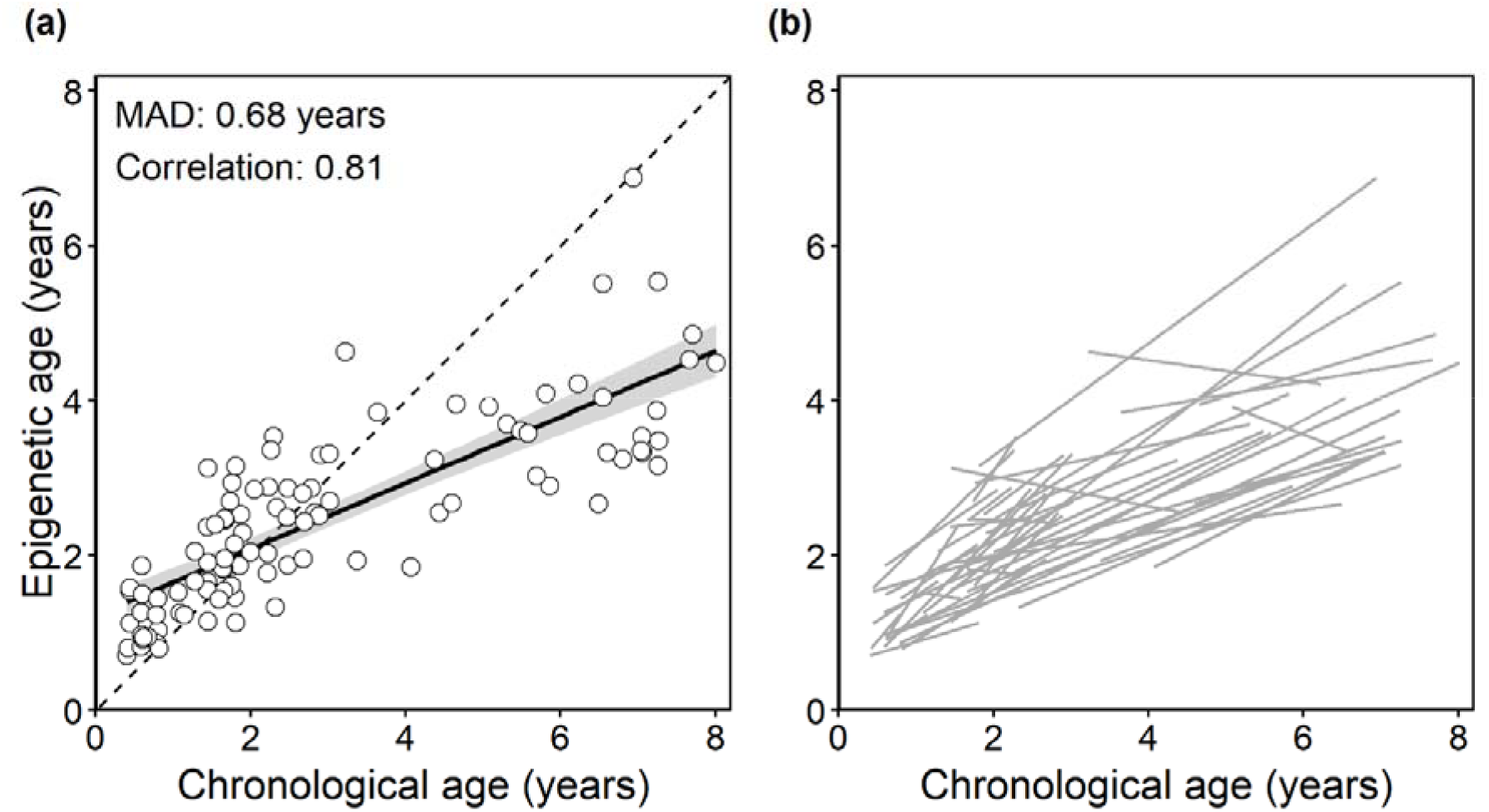
Epigenetic age in relation to chronological age, with epigenetic age predicted using a leave-one-individual-out cross-validation (LOIOCV) approach on 100 zebra finch samples and trained on logarithmic transformation of age. (a) Regression over all individual samples with the solid line representing the fitted linear regression and the shaded area being the 95% CI. Dashed line shown as reference representing the expected relation if chronological and predicted age were identical (Y=X). (b) Spaghetti plot showing within-individual change in epigenetic age in relation to chronological age. Each grey line represents one individual.

Epigenetic clocks can be constructed using a variety of modeling and validation strategies. We trained our clock using a LOIOCV approach, where a clock is trained on all individuals except one, then tested on the left-out individual. An alternative approach using an 80/20 train–test split yielded a very similar performance, achieving a MAD of 0.78 years and a correlation of r⍰=⍰0.72 (*p* < 0.001) between predicted and chronological age. This clock shared 27 of its 53 clock CpGs with the LOIOCV-trained clock.

To determine whether genomic functional regions were depleted, enriched or neither in selected clock CpGs relative to the total number of CpGs after filtering, we performed Fisher’s exact tests per region. Clock CpGs were mostly located in introns (42.9%) and intergenic regions (41.2%), followed by exons (10.1%) and promoters (5.8%; Fig.3b). Binomial tests comparing the observed and expected number of clock CpGs in each genomic category indicated that numbers in none of the categories deviated significantly from chance expectation (promoters: *p* = 0.096, exons: *p* = 0.58, introns: *p* = 0.36, intergenic regions: *p* = 0.45). The clock CpGs were located in or near 101 unique genes, 40 of which were in LOC genes with uncharacterized function.

Correlations between DNAm and age at the 119 CpG sites included in the clock ranged from - 0.567 to 0.573, with trends being positive at 39 sites, and negative at 80 sites. The absolute elastic net coefficient of clock CpGs was significantly positively correlated with their absolute correlation between DNAm and age (Fig.S5; r=0.37, *p* < 0.001). However, as illustrated by the scatter plot, the clock also includes CpGs with moderate correlations with age, indicating that the elastic-net selection favors sets of CpGs with joint predictive value and complementary information, not necessarily the sites with highest correlation with age.

Age may not affect the sexes equally (e.g. Briga et al. 2017), and we earlier found rearing brood size effects on DNAm to be strongly sex-specific (Tangili, Palsbøll, et al., 2026). We therefore explored sex dependence of age-related DNAm by making sex-specific epigenetic clocks, using data from either males or females only, while only using CpG sites shared by all samples. Both clocks predicted chronological age with somewhat lower accuracy than the clock developed using all data, as expected given the reduced sample size (female only clock: r=0.61, MAD = 0.91 years, Fig.S5a; male only clock: r = 0.74, MAD = 0.82 years, Fig.S5b). Interestingly, the number of CpG sites included differed strongly between the clocks, with 29 CpG sites in the females only clock, and 161 CpG sites in the males only clock (Table S1). Distribution of clock CpG sites over genomic location was very similar for the two sexes (respectively for females and males: promoters: 10 / 5%; exons: 10 / 12%; introns: 45 / 43%; intergenic: 34 / 40%; Fig.S5c).

Eighteen out of 29 CpG sites identified in the EWAS were shared with the clock for chronological age (Fig.S4, Fig.S5d). This number reduced to four (14%) between the EWAS and female clock and to 17 (11%) between the EWAS and male clock (Fig.S5d). Three sites that were shared between the EWAS and original clock were all located in the promoter of the *SOCS2* gene, and DNAm decreased significantly with age at all three sites (Fig.S6).

### Epigenetic age acceleration and phenotype

As expected, since samples were balanced for sex and brood size across the age distribution, neither rearing brood size nor sex was significantly associated with epigenetic age (Table S3, Fig.S7). However, birds raised in large broods had significantly higher age acceleration than birds raised in small broods (Table 2, Fig.5a), in line with their shorter life expectancy (Briga et al. 2017). Sex-specific post-hoc tests revealed that the rearing brood size effect was sex-limited, with significant epigenetic age acceleration in females reared in a large broods, compared to females reared in small broods (estimate = 0.19, t_(22)_ = 2.27, *p* = 0.03). There was no such effect in males (estimate = -0.02, t_(24)_ = -0.23, *p* = 0.82), but note that the interaction between sex and brood size did not quite reach statistical significance. An alternative way to examine early-life effects is to use early life growth rate, which was affected by the brood size manipulation (Briga et al., 2017; Tangili, Briga, et al., 2026). Epigenetic age acceleration of the samples taken early in adult life was negatively correlated with early-life growth rate (r = -0.32, *p* =0.04; Fig.5b), indicating that birds that grew less in early life were epigenetically older in adulthood. This pattern was independent of sex (growth rate ^*^ sex : *p* = 0.17). In contrast, epigenetic age acceleration of the samples taken late in adult life was not significantly correlated with early-life growth rate (r = -0.07, *p* = 0.64). The interaction between early-life growth rate and age at sampling (early versus late in life) was close to statistical significance (*p*=0.057), suggesting that the effects of early-life growth on epigenetic age acceleration diminish with age.

**Figure 5.**
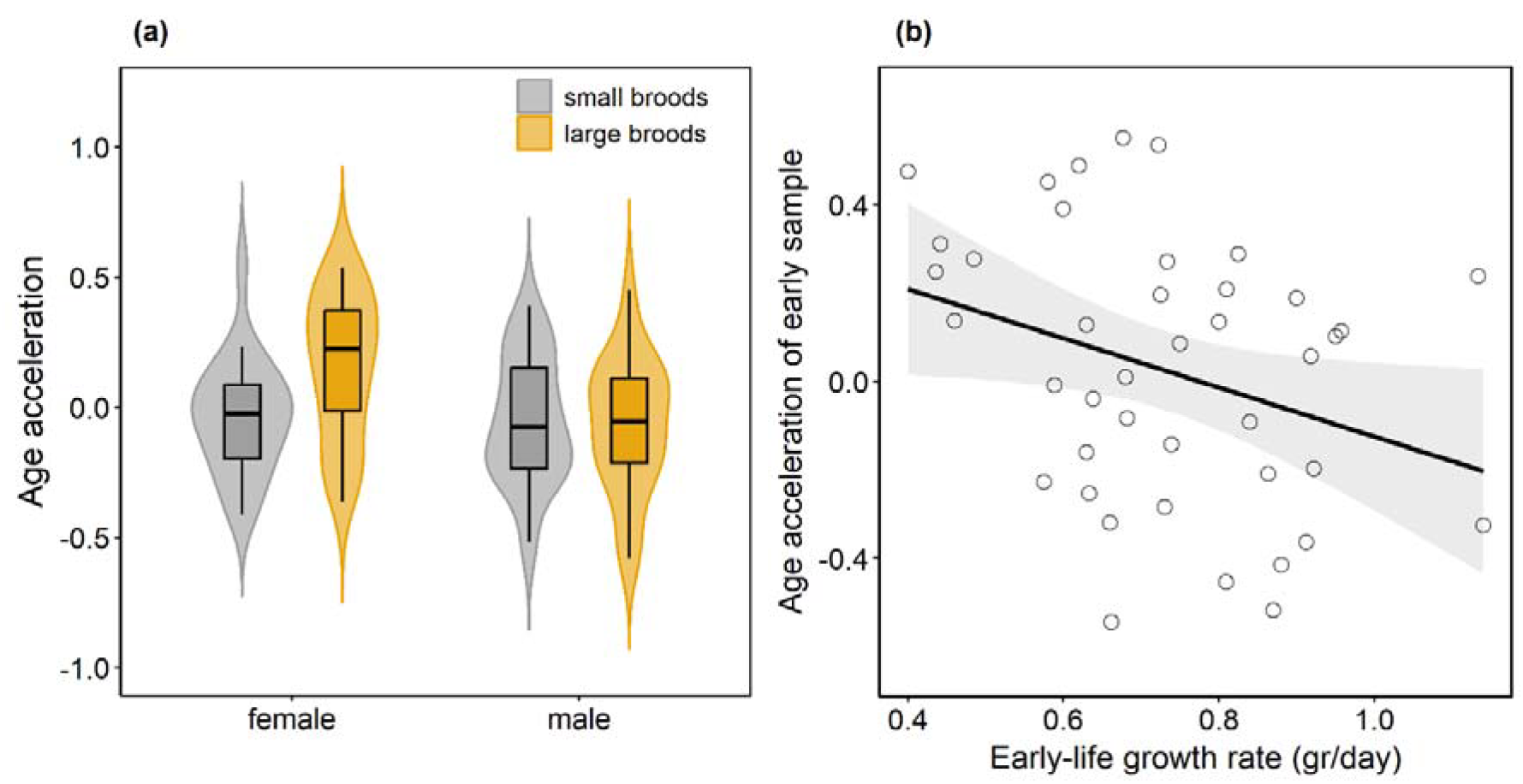
Epigenetic age acceleration in relation to sex and early-life conditions. Age acceleration estimates are residuals (N=100) of the linear mixed effects model of chronological vs. epigenetic age with individual id as a random effect from the epigenetic clock trained on logarithmic transformation of age. (a) Age acceleration in relation age sex and rearing brood size. (b) Age acceleration in relation to early-life growth rate (until day 15, N=50) of the early sample per individual. Solid line represents the fitted linear regression and the shaded areas are the 95% CI.

**Table 2.** Results of linear mixed effects model on the effects of brood size and sex on epigenetic age acceleration (calculated as the residuals of the linear mixed effects model of chronological vs. epigenetic age with individual id as a random effect) from the epigenetic clock trained on logarithmic transformation of age. Reference categories are small broods and females.

|  | Residuals (age acceleration) |  |  |  |
| --- | --- | --- | --- | --- |
| | Estimate (± SE) | t-value | df | Pr(>\|t\|) |
| Fixed effects |  |  |  |  |
| Intercept | -0.04 (± 0.06) | -0.59 | 46 | 0.56 |
| brood size | **0.19 (± 0.09)** | **2.21** | **46** | **0.03 \*** |
| sex | -0.01 (± 0.09) | -0.17 | 46 | 0.87 |
| brood size * sex | -0.21 (± 0.12) | -1.76 | 46 | 0.09 |
| Random effects |  |  |  |  |
| bird id | 0.03 |  |  |  |
| Residual | 0.04 |  |  |  |

We previously developed a Supervised Machine Learning methylation index (SMLmi), that accurately predicted rearing brood size for 89% of the samples utilizing the DNAm of 455 CpG sites (Tangili, Palsbøll, et al., 2026). We therefore predicted a positive correlation between the rearing brood size index and the age acceleration we estimated in the present paper, especially for females we found to have significant epigenetic age acceleration when raised in large broods. Contrary to this prediction, we found no significant correlation between the SMLmi and epigenetic age acceleration for either sex (both *p*>0.05, Fig.S8, Table S3), which shows that individuals that have high epigenetic age acceleration do not show an epigenetic state that is correlated with being reared in a large brood.

### Lifespan prediction

Our ultimate aim was to test whether a high epigenetic age relative to chronological age predicted a reduced life expectancy, as robustly demonstrated in humans (Levine et al., 2018; Lu et al., 2019). We tested this hypothesis firstly by analyzing epigenetic age in relation to age and lifespan, but found no association, neither as main effect, nor in interaction (Table S4). We then related lifespan to age acceleration estimated for the early samples (i.e. the first sample in adult life of each individual) and found that this also did not significantly predict lifespan (Fig.6a; estimate = 0.39, t_(48)_=0.33, *p* = 0.73) or remaining lifespan (Fig.6b; estimate = 0.3, t_(48)_=0.29, *p* = 0.77). Neither did epigenetic age in early life predict remaining lifespan (Fig.6c; estimate = 0.01, t_(48)_=0.05, *p* = 0.96). Interpretations of the relationship between epigenetic age and lifespan are constrained by collinearity; the clock was trained on chronological age, and chronological age was strongly associated with lifespan as the individuals which lived longer were also the ones which were sampled later in life. Therefore, under these circumstances it is difficult to isolate the independent predictive contribution of epigenetic age to lifespan variation.

**Figure 6.**
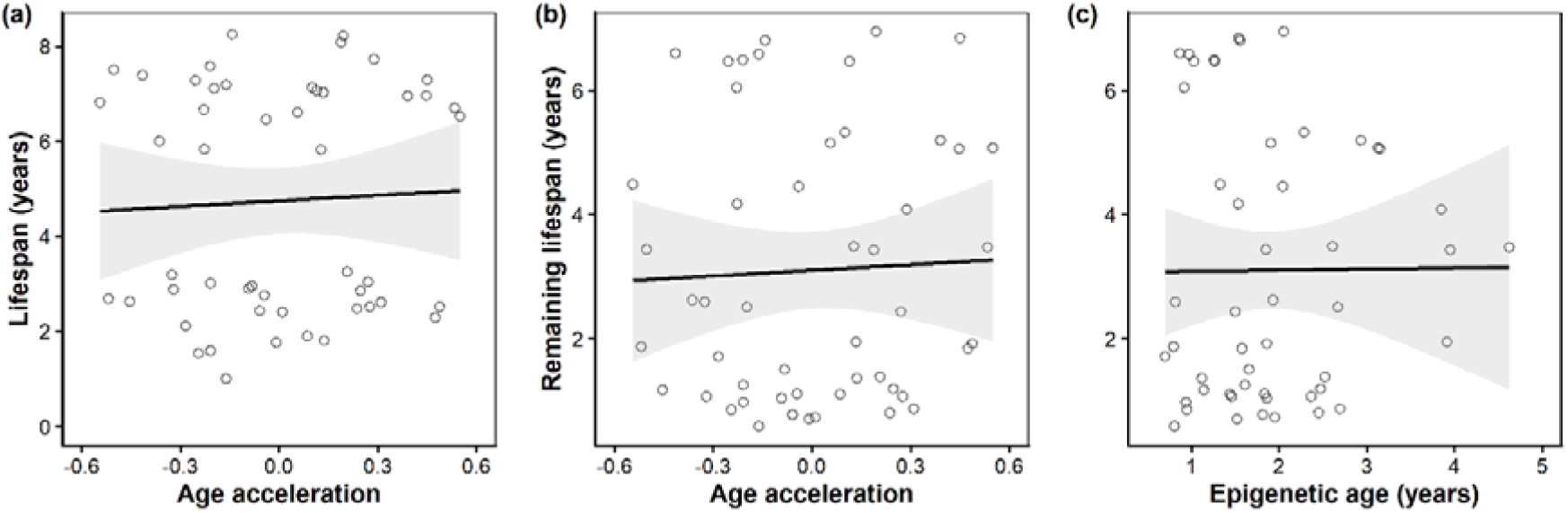
(a) Epigenetic age acceleration (residuals of the linear model of chronological in relation to lifespan (a) and remaining lifespan (b) for the early samples. (c) Predicted epigenetic age in relation to remaining lifespan. Solid lines represent the fitted linear regressions and the shaded areas are the 95% CI.

If epigenetic aging reflects (aspects of) biological aging, we would expect a significant relationship between epigenetic aging rate and lifespan. Consistent with this expectation, utilizing our longitudinal design, we find that short-lived individuals have a higher rate of epigenetic aging than long-lived individuals (Fig.7, Table 3).

**Figure 7.**
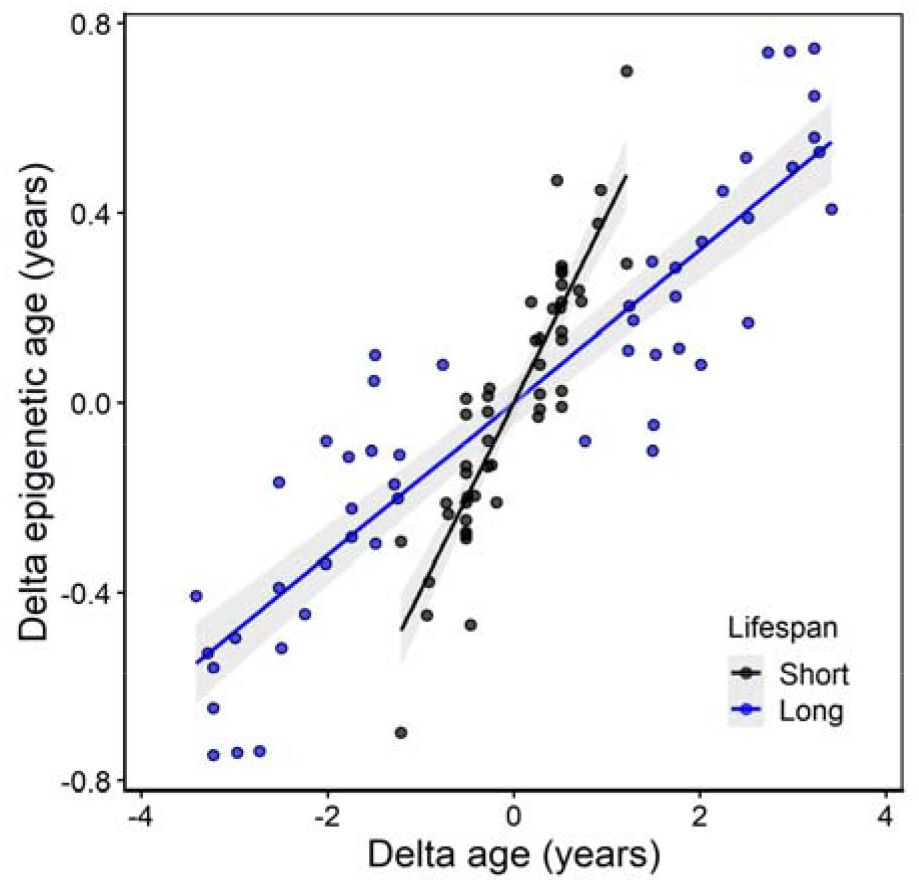
Individual epigenetic aging rate and lifespan. Relationship between the within-individual change in epigenetic age and delta age per bird (delta age = age at sampling – mean of the two sampling ages per individual). Different colors denote individuals with short vs. long lifespans, the solid lines represent the fitted linear regression and the shaded areas are the 95% CI. Individuals with “short” lifespans had an average lifespan of 2.44 ± 0.57 years and individuals with “long” lifespans had an average lifespan of 7.04 ± 0.65 years. Note that grouping in short and long lifespan is for graphical purposes only – the statistical analysis was based on lifespan as a continuous variable (Table 3).

**Table 3.** Results of linear mixed effects model on the effects of individual epigenetic aging rate and lifespan. All age variables are expressed in years.

|  | Epigenetic age (years) |  |  |  |
| --- | --- | --- | --- | --- |
| | Estimate (± SE) | t-value | df | Pr(>\|t\|) |
| Fixed effects |  |  |  |  |
| Intercept | **6.197(±0.09)** | **72.18** | **47** | **<0.001 \*\*\*** |
| delta age | **0.437(±0.07)** | **5.95** | **48** | **<0.001 \*\*\*** |
| average age | **0.199(±0.06)** | **3.26** | **47** | **0.002 \*\*\*** |
| lifespan | -0.019(±0.04) | -0.51 | 47 | 0.613 |
| lifespan*delta age | **-0.038(±0.01)** | **-3.61** | **48** | **<0.001 \*\*\*** |
| Random effects |  |  |  |  |
| bird id | 0.05 |  |  |  |
| Residual | 0.05 |  |  |  |

### Doom clock

As complement to the analyses of the relationship between age acceleration and lifespan (Fig.7) and individual epigenetic aging rate and lifespan (Fig.8), we trained an epigenetic clock on realized remaining lifespan. Due to our selection of samples early and late in adulthood of each individual, the variation in remaining lifespan after the late samples was too low to be informative (SD=0.56 years). We therefore trained an epigenetic clock on the remaining lifespan using the early samples only (N = 50; mean±SD remaining lifespan: 3.1±2.16 years). This ‘doom clock’ significantly predicted remaining lifespan (r = 0.37, p = 0.009; Fig.8), with a MAD of 1.75 years. The doom clock was based on DNAm of only eight CpG sites, substantially less than used in the clock of chronological age (n=119 CpG sites), and none of the eight CpG sites was shared with the CpGs of the chronological age clock.

**Figure 8.**
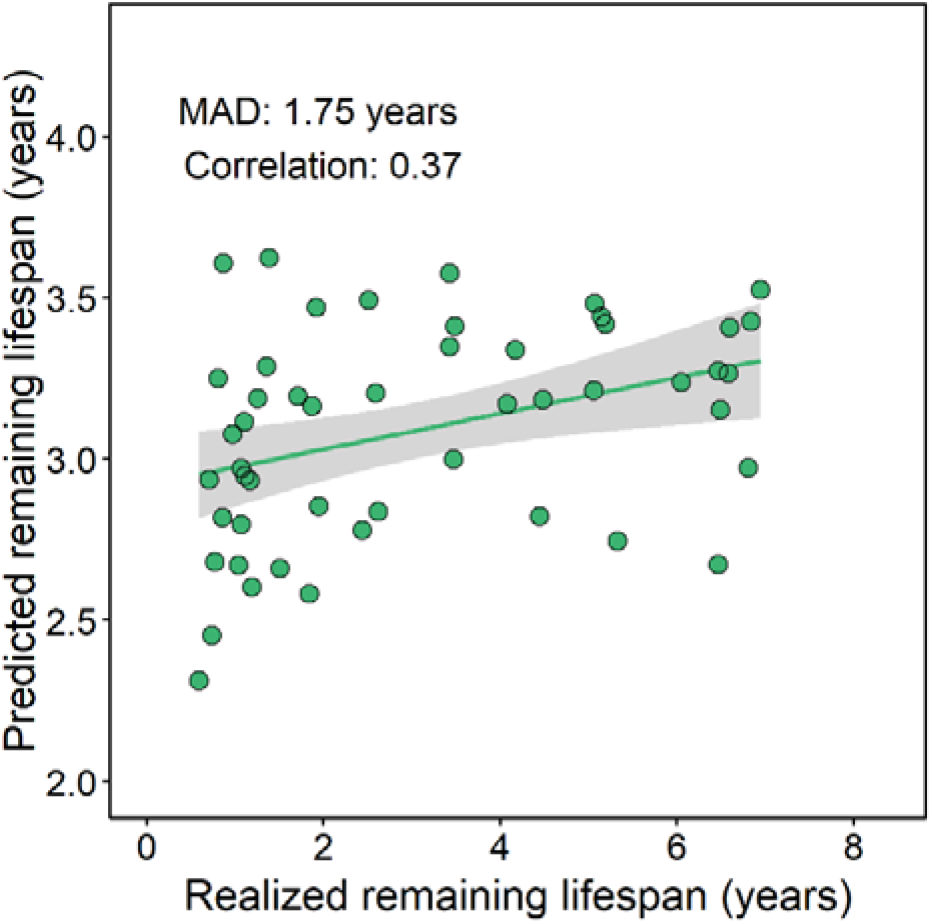
Predicted remaining lifespan in relation to realized remaining lifespan, predicted using a leave-one-out cross-validation (LOOCV) approach on only the early sample taken from each of the 50 zebra finches. Solid line represents the fitted linear regression and the shaded area is the 95% CI. Note that the two axes are not on the same scale.

## Discussion

We investigated associations between age and DNAm in three different ways, on a genome-wide level, through an epigenome wide association study (EWAS), and through the development of an epigenetic clock of age, with the overarching aim to provide insight on associations between DNAm and individual characteristics, including lifespan.

Genome-wide levels of DNAm decrease with age across mammalian tissues (Berdyshev et al., 1967; Vanyushin et al., 1973; Wilson et al., 1987), a pattern that has been suggested to contribute to the aging process (Unnikrishnan et al., 2018). In humans, hypomethylation with age is concentrated in repetitive genomic sequences where the gene expression leads to an aging phenotype (Bollati et al., 2009). Here we report genome-wide hypomethylation with age in the zebra finch, independent of sex. Hypomethylation was previously reported in two other bird species; chicken hens (Gryzinska et al., 2013) and female common terns (but not males, *Sterna hirundo, Meyer et al., 2023*), suggesting hypomethylation with age may be a common pattern in birds. Genome-wide levels of DNAm were not correlated to brood size, contrasting previous findings revealing a positive correlation at 107 CpG sites between DNAm and natal brood size in zebra finches (Sheldon et al., 2018). Importantly, we found that autosomal DNAm declined faster with age in short-lived individuals compared to longer-lived individuals, consistent with the hypothesis that hypomethylation may be linked to lifespan (Xiao et al., 2016).

The genome-wide hypomethylation trend was also observed on the individual CpG site level, with DNAm decreasing with age across the majority of CpG sites. This was confirmed in the EWAS of chronological age, which identified twenty-nine CpG sites where DNAm correlated significantly with chronological age, with DNAm decreasing with age at 83% of these sites. Promoters were significantly enriched and introns significantly depleted for EWAS significant CpGs. The number of CpG sites with significant correlations between DNAm and age emerging from the EWAS was substantially lower than identified earlier in the same population, with no overlap between the two CpG sets (Tangili et al., 2025). We attribute this discrepancy to a methodological difference: the earlier analysis was longitudinal, correlating DNAm with within-individual age variation, while the EWAS was predominantly based on cross-sectional data. The different number of CpG sites identified and absence of overlap between the two sets illustrates the value of a longitudinal approach.

Following the EWAS, we developed an epigenetic clock of age that predicted chronological age with high accuracy, with the MAD being equivalent to 8.2% of the maximum lifespan in our dataset (8.25 years). This MAD scaled to maximum lifespan was lower than the scaled MAD of earlier avian epigenetic clocks (short-tailed shearwater, *Ardenna tenuirostris* : 13.3%, De Paoli-Iseppi et al., 2019; great tit : 13%, Haller et al., 2025; chestnut-crowned babbler, *Pomatostomus ruficeps* 8.4%, Gerber, Schrey, et al., 2025). We previously found CpG sites with age-related DNAm to be concentrated on the Z chromosome (Tangili et al., 2025), but none of those were retained in the epigenetic clock. Apparently, robust age prediction relies primarily on autosomal DNAm. While there was some overlap between the CpG sites identified in the EWAS of age and the clock CpGs, the clock CpGs also included sites with moderate correlations between DNAm and age. Thus, the epigenetic clock utilized both CpGs that are robustly age-associated and additional sites that contribute to prediction through multivariate weighting rather than by having the strongest univariate correlations (Higgins-Chen et al., 2022; A. Li et al., 2022).

We trained our clock using a LOIOCV approach, but using an 80/20 train–test split yielded a very similar performance; this clock shared 27 of its 53 clock CpGs with the LOIOCV-trained clock. These results indicate robustness of our epigenetic clock to differences in training and testing design with respect to the attained accuracy, but less so with respect to the included CpG sites. The zebra finch clock showed a typical regression of the mean pattern (Bland & Altman, 1994; Simpson & Chandra, 2021). When the purpose if the main aim of clock is to predict an unknown chronological age, e.g. in the context of conservation or forensics, prediction accuracy can be further increased by applying a bespoke transformation to the clock output.

Sex and early-life adversity (nutritional stress and sibling competition) impacted lifespan in our study population (Briga et al., 2017), and we therefore tested for effects of these factors on epigenetic age acceleration. Overall, age acceleration did not differ between sexes or between birds experiencing different levels of sibling competition. There was however some evidence for an interaction, with only females that experienced more sibling competition having higher epigenetic age acceleration, indicating they were biologically older. That this effect was sex limited is consistent with evidence that female zebra finches are more vulnerable to early life food restriction than males (Fernandes Martins, 2004). High growth rate of nestling birds is generally associated with high fitness prospects (Gerritsma et al., 2022) and, in line with this pattern, we found a negative association between epigenetic age acceleration and early life growth. An interesting aspect of this finding was that it only applied to the samples taken in early adulthood, suggesting that this signature of early life adversity fades with age.

Humans with a higher epigenetic age than expected for their chronological age have a shorter life expectancy (Kuo et al., 2026; Levine et al., 2018; Lu et al., 2019), but we could not confirm this in our study, at least in a cross-sectional analysis. In contrast, a longitudinal analysis of epigenetic aging rate revealed a significant association with lifespan. Thus, in our study it was the rate of change of epigenetic age within individuals that predicted lifespan rather than the absolute level. In line with our findings, available longitudinal human studies also observed the rate at which epigenetic age increased with time to predict mortality risk (Drewelies et al., 2025; Kuo et al., 2026). These and our findings underscore the importance of longitudinal sampling in studies of (epigenetic) aging.

Having established clear associations of DNAm with aging and lifespan, we asked whether DNAm could also be used to predict *remaining* lifespan directly. Using only early-life samples, we trained an epigenetic “doom clock” that significantly predicted remaining lifespan, although the precision was modest. None of the eight CpGs used in the doom clock were featured in the chronological age clock, suggesting that CpG sites that predict remaining lifespan and chronological age might capture partly distinct biological processes. Remaining-lifespan clocks can provide a potentially powerful biomarker to identify variation in longevity prospects early in life. ‘Doom’ clocks may have clinical relevance, creating opportunities to identify and then address vulnerabilities early in life. It is important to consider that although epigenetic age acceleration and epigenetic clocks trained directly on survival-related outcomes such as the doom clock are both informative, they represent distinct constructs. Epigenetic clocks trained directly on individual-specific outcomes are expected to be more precise in predicting those outcomes compared to indirect measures of age acceleration derived by epigenetic clocks trained on chronological age.

An open question is whether DNAm at clock CpGs functionally contributes to the aging process (Horvath & Raj, 2018). Five of the clock CpGs we identified were located in the Suppressor of cytokine signaling 2 (SOCS2) gene, four in the promoter and one in an intron. The four clock CpGs in the promoter of this gene all lost DNAm with age and were also identified as significant CpGs in the EWAS of age, while two of them were shared by all age-related DNAm analyses performed in this study. SOCS2 is a key negative regulator of cytokine signaling and through the Janus kinase and signal transducer and activator of the transcription (JAK-STAT) pathway is known to mediate inflammation, growth hormone responses and immune homeostasis while its aberrant regulation has been linked to both inflammatory and neoplastic diseases (Letellier & Haan, 2016). Previous research has also shown that absence of SOCS2 expression leads to reduced lifespan in mice which indicates its potential involvement of this gene in regulating aging, potentially through its impact on plasma Insulin-like Growth Factor 1 (IGF1) concentration (Casellas & Medrano, 2008). The hypomethylation of the SOCS2 promoter with age likely upregulates its expression and contributes to age-related changes in growth, tissue maintenance, and immune function which can in turn affect longevity. Other genes in or near clock CpGs include the *IGFBP3* gene (Insulin-like Growth Factor Binding Protein 3) which regulates IGF1 activity and inhibits cell proliferation by inhibiting telomerase activity (Kwon et al., 2023), the *RUNX2* gene (Runt-related transcription factor 2) whose hypomethylation is associated with osteoporosis and age-related dysfunctional calcification in humans (Yalaev et al., 2024) and the *YAP1* gene (Yes-associated protein 1), whose activity declines with age and leads to cellular senescence and impaired tissue regeneration (Elster & von Eyss, 2020). One of the doom clock CpGs is located on an intron in the ETS Variant Transcription Factor 1 (*ETV1*) gene, part of the E-twenty-six transcription factor family, which has been shown to determine longevity in multiple tissues and animal taxa (Dobson et al., 2019). Although our findings suggest that age-related change in DNAm of clock CpGs may be functionally linked to the aging process, expression analyses connecting DNAm to gene expression and the aging phenotype will be required to confirm this relationship.

Epigenetic marks have been previously found to predict mammalian species’ maximum lifespan and other life-history traits (Horvath et al., 2022; C. Z. Li et al., 2024), yet individual lifespan prediction using epigenetic clocks has been limited to humans and mice (Levine et al., 2018; Lu et al., 2019; Thompson et al., 2018). The predictive ability of our doom clock was found to be limited but non-random as reflected by a significant, positive correlation between realized and predicted remaining lifespan. This finding suggests that epigenetic marks contain biologically meaningful information about future survival, while also underscoring that additional environmental, genetic, and stochastic factors likely contribute to variation in remaining lifespan. Although our doom clock serves as a first proof-of-concept, it is evident that larger datasets would be needed to confirm our findings and more robustly test these hypotheses.

## Conclusion

We have here developed the first epigenetic clock of age for the zebra finch, a model avian species, which achieved high predictive accuracy. Several of the CpG sites used for the clock of chronological age were found to be located in or near genes involved in development, aging and longevity. Critically, we have connected the epigenetic clock output to the phenotype; females raised in large broods, which are known to have shorter lifespans, showed increased epigenetic age acceleration. Moreover, individuals which grew less during the nestling stage were epigenetically older, at least early in adult life. Importantly, individuals with higher within-individual epigenetic aging rate had shorter lifespans, which mirrors the pattern we have previously found for telomere length in the same zebra finch cohort, with telomere shortening rate but not absolute length predicting lifespan (Tangili, Mulder, et al., 2026). A doom clock predicted remaining lifespan with moderate accuracy utilizing the DNAm of a set of CpGs that did not overlap with the clock CpGs of chronological age, pointing to the two clocks capturing distinct aging processes. Together, our findings extend the promise of remaining lifespan prediction using DNAm information beyond mammalian species. We hope that this study encourages researchers to include longitudinal sampling whenever possible and utilize epigenetic clocks beyond chronological age estimation, as studies like the present hold the potential to uncover the exact molecular mechanisms that dictate individual lifespan variation as well as the ones through which the environment shapes life-history tradeoffs.

## Acknowledgements

We thank the Center for Information Technology at the University of Groningen for support and access to the Hábrók high-performance computing clusters. We express our gratitude to Ellis Mulder for her help in the laboratory and to Michael Briga and Blanca Jimeno for running the long-term experiment. We are also indebted to the animal caretakers at the University of Groningen as well as numerous students whose invaluable help made this project possible.

## Ethics approval

All methods and experiments detailed in this manuscript were performed under the approval of the Central Committee for Animal Experiments (Centrale Commissie Dierproeven) of the Netherlands, under licenses AVD1050020174344 and AVD1050020184967.

## Author contributions

MT and SV conceived and designed the study. MT performed the bioinformatics and data analysis under the supervision of SV and PJP. MT and SV wrote the manuscript with input from all authors. All authors approved the final version of the manuscript.

## Data availability

Raw reads of the whole methylomes of all samples are available at the National Center for Biotechnology Information under BioProject ID PRJNA1108628. The bioinformatics code for methylation extraction is available in Tangili et al. (2024) and R scripts and additional data are available in <u>10.6084/m9.figshare.33514576</u>.

## Supplementary information

**Figure S1.**
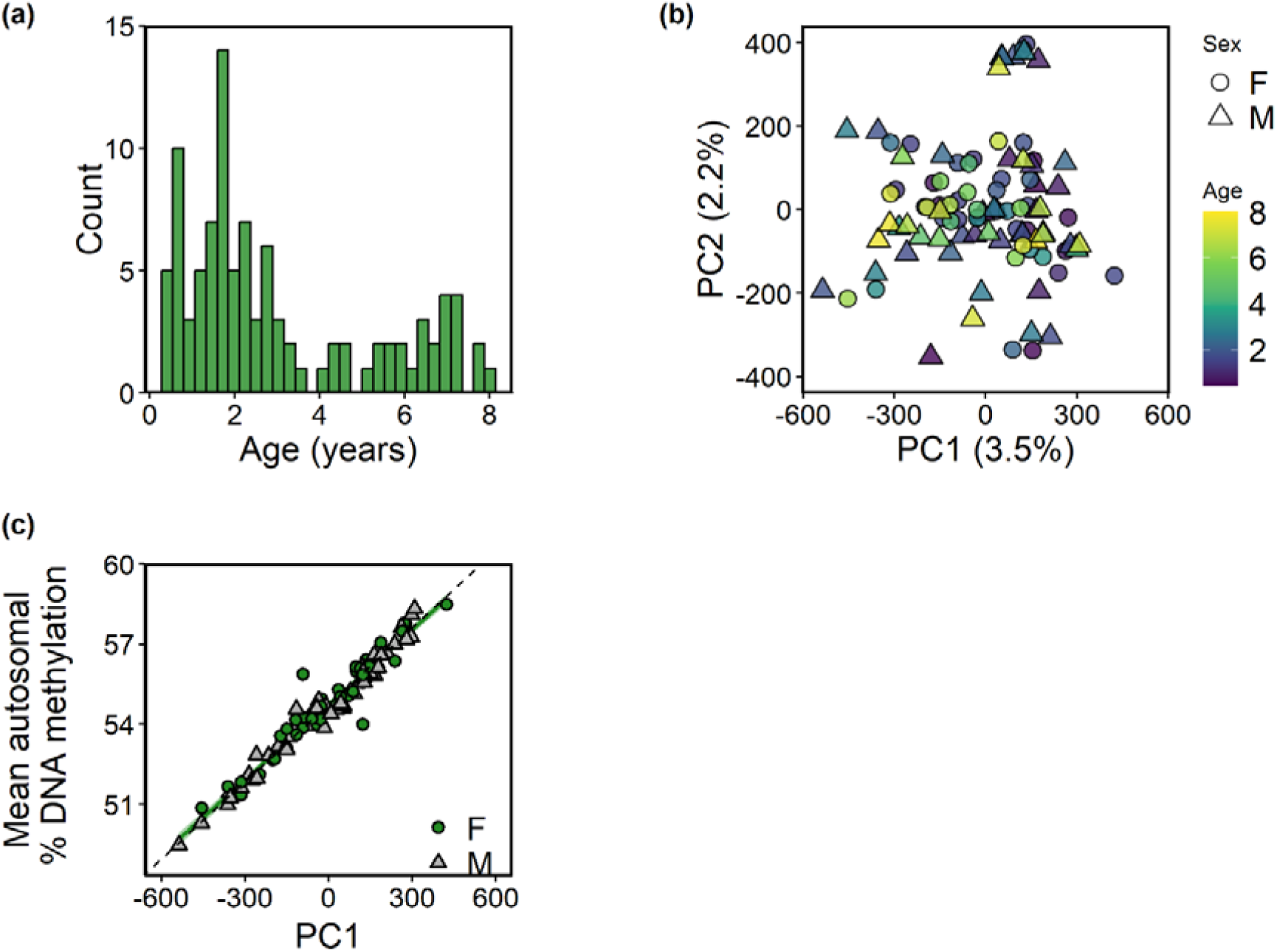
(a) Age at sampling distribution of the dataset (N=100 samples from 50 adult individuals). (b) Principal component analysis based on the methylation of the CpG sites that passed filtering. Color indicates the age and shape the sex of each sample. (c) The first principal component (PC1) in relation to mean autosomal % DNA methylation per sample.

**Figure S2.**
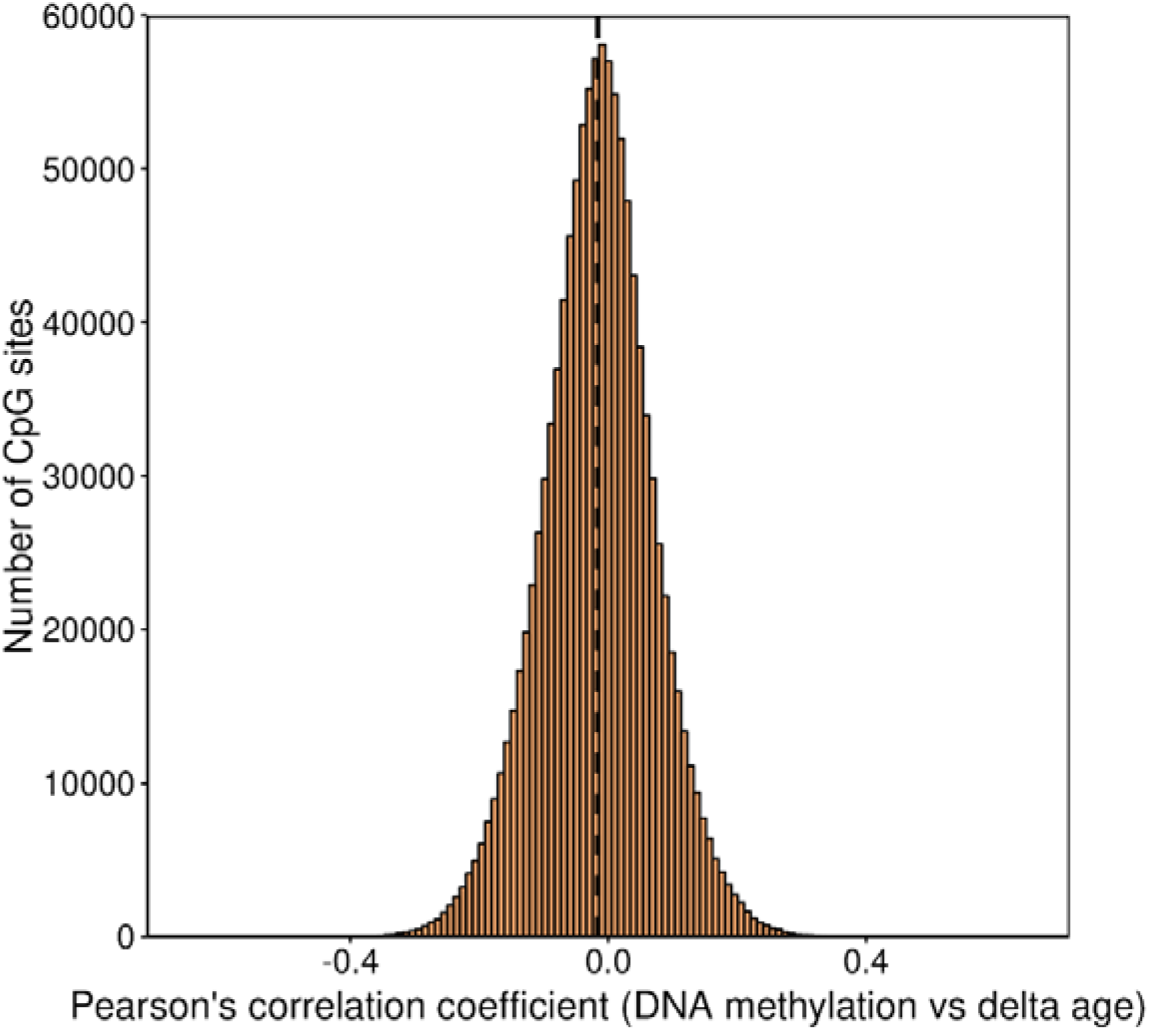
Distribution of Pearson correlation coefficients between delta age (chronological age-average age per individual) and DNA methylation for all CpG sites that passed filtering. The dashed vertical line denotes the mean (-0.016, 95% CI = -0.017 -0.016)

**Figure S3.**
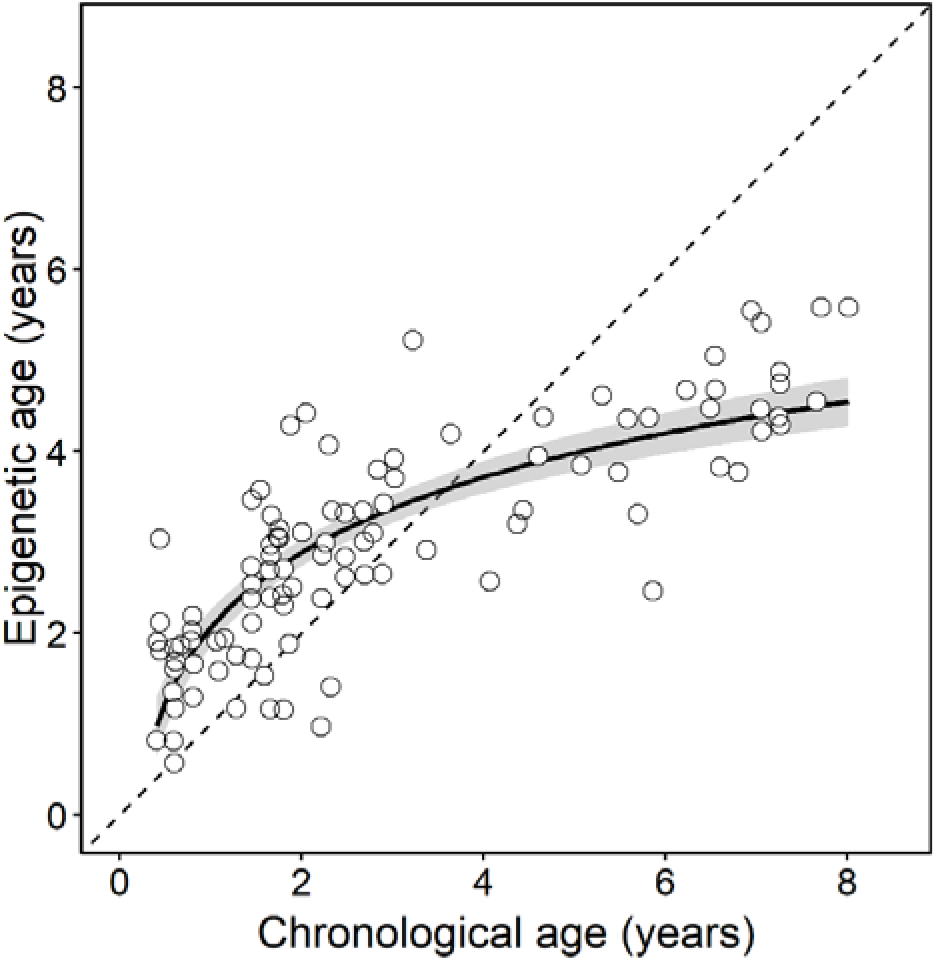
Relationship of chronological age and epigenetic age predicted using a leave-one-individual-out cross-validation (LOIOCV) approach or 100 zebra finch samples. The dotted line represents the regression line if chronological and predicted age were identical (Y=X). Solid line represents the fitted logarithmic regression of epigenetic age on chronological age and the colored area is 95% CI.

**Figure S4.**
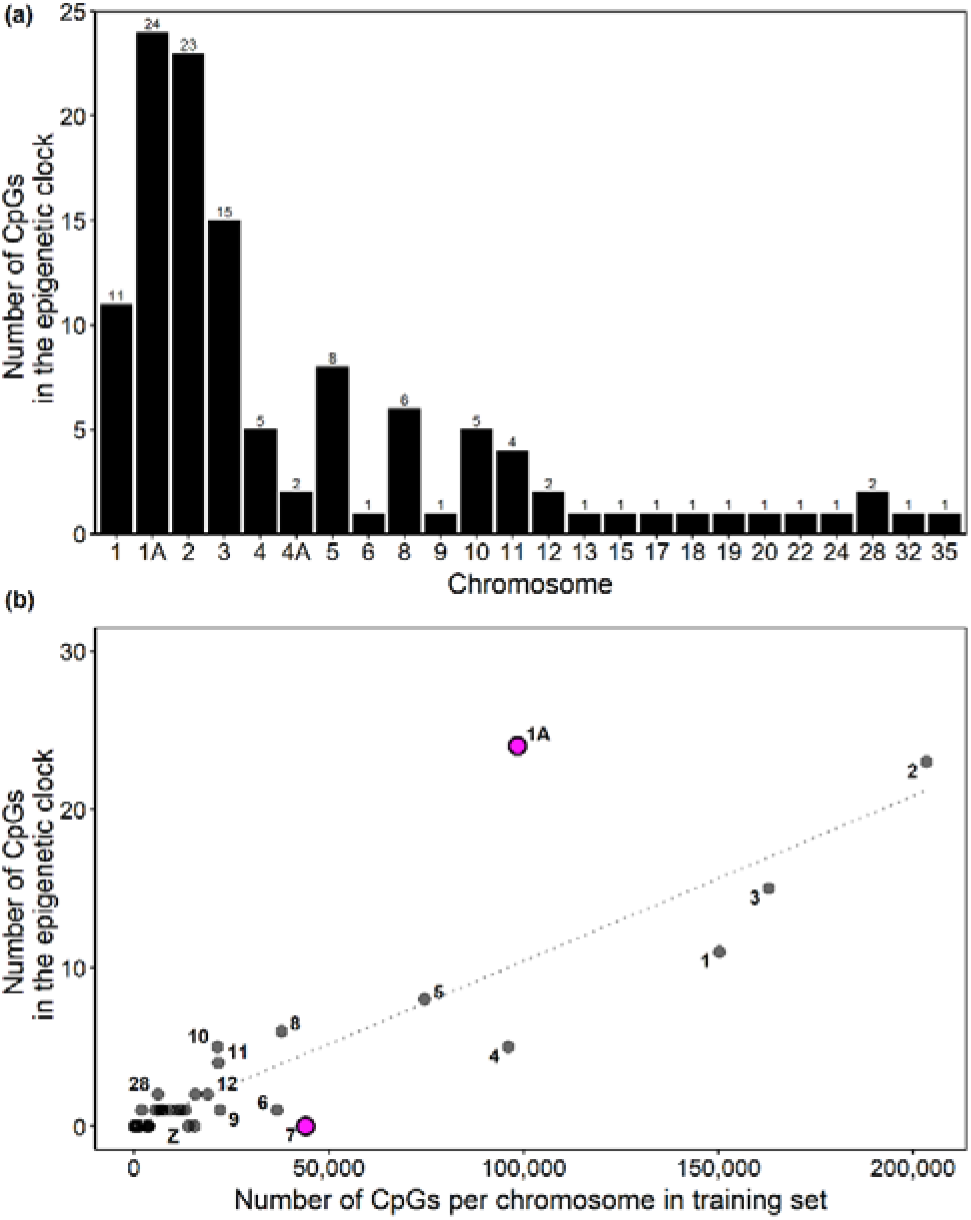
(a) Distribution of CpG sites in the final epigenetic clock model over chromosomes. (b) Number of epigenetic clock CpG sites in relation to total CpG sites per chromosome in the training dataset. The dotted line represents null expectation, if clock CpGs were equally distributed among chromosomes. Chromosome 1A was significantly enriched and chromosome 7 significantly depleted for clock CpGs.

**Figure S5.**
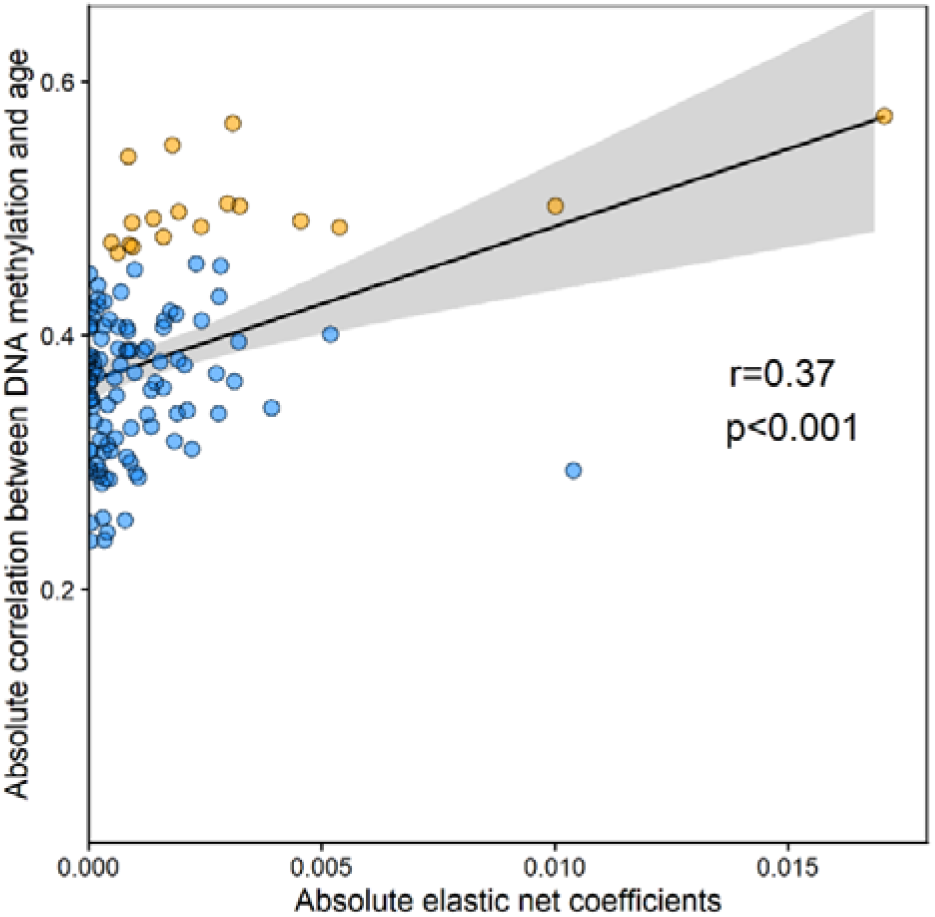
Relationship between absolute elastic net coefficients and absolute correlation between DNA methylation and age of clock CpGs. Orange points represent the CpG sites shared between the epigenetic clock and the EWAS of chronological age. Solid line represents the fitted linear regression and the shaded area is the 95% CI.

**Figure S5.**
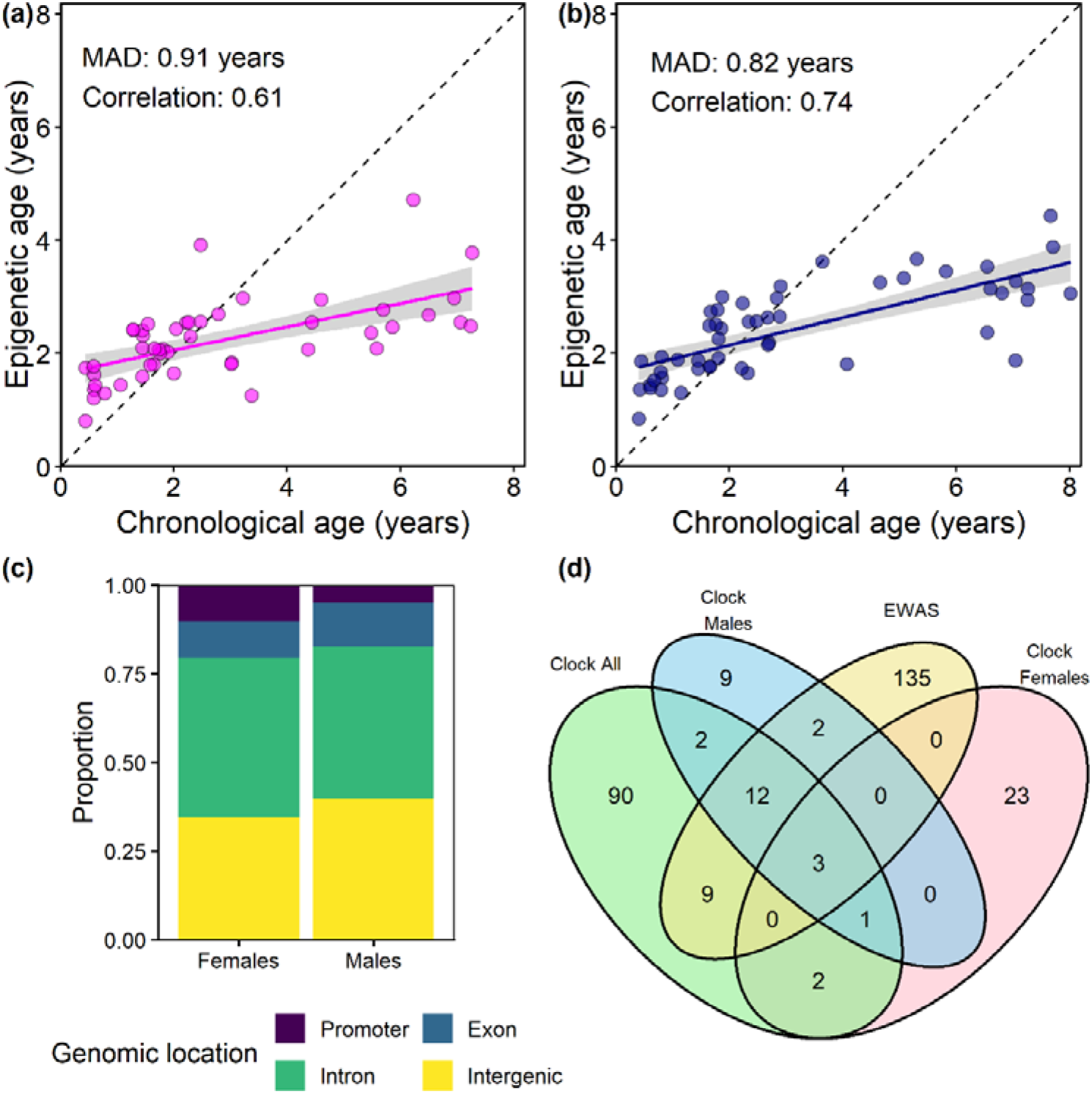
Female (a) and male-specific (b) epigenetic clocks. (c) Genomic location of clock CpGs in the female and male-specific clocks. (d) CpG sites shared among the EWAS of chronological age, the clock including all samples, the female and male-specific clocks.

**Figure S6.**
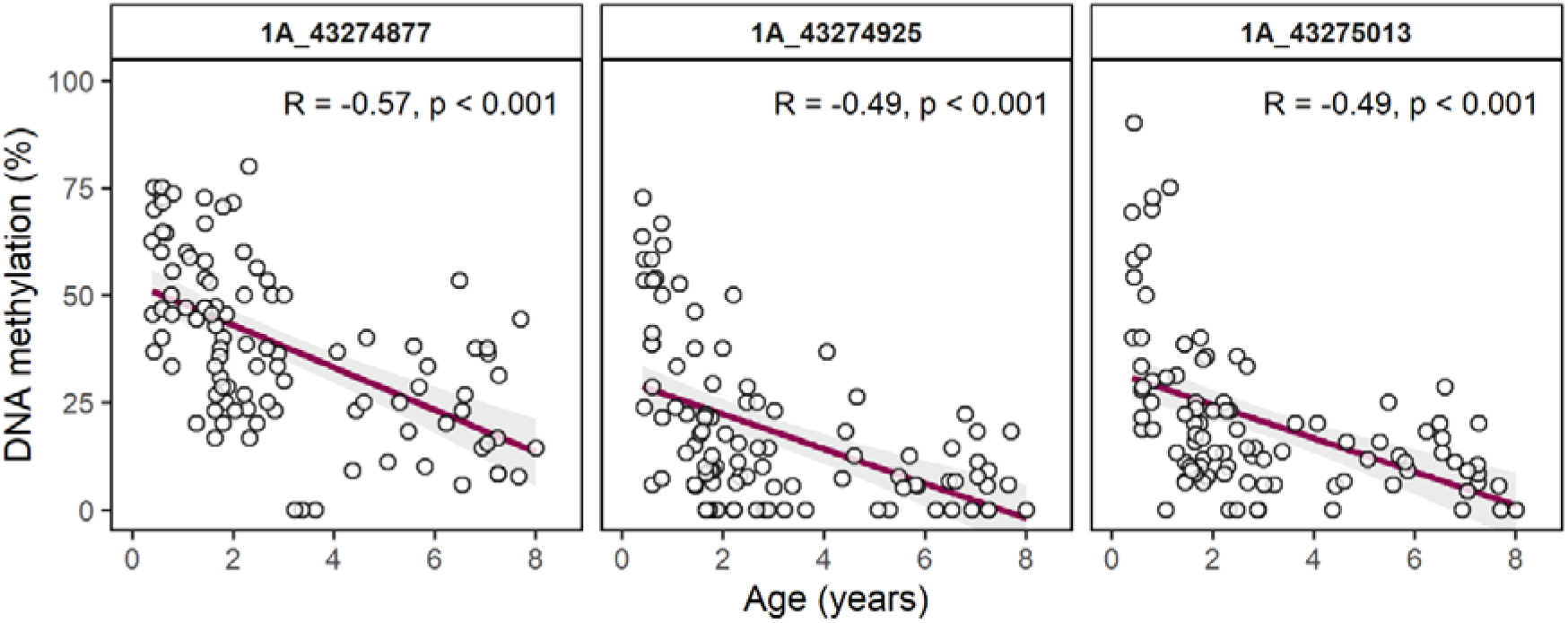
Age-related decline in the methylation of the three CpG sites on the promoter of the SOCS2 gene which were shared by the EWAS of chronological age and the clock CpGs. 1A_43274877 and 1A_43274925 were shared among the EWAS of chronological age and all the epigenetic clocks. Solid lines represent the fitted linear regressions, and the shaded areas are the 95% CI.

**Figure S7.**
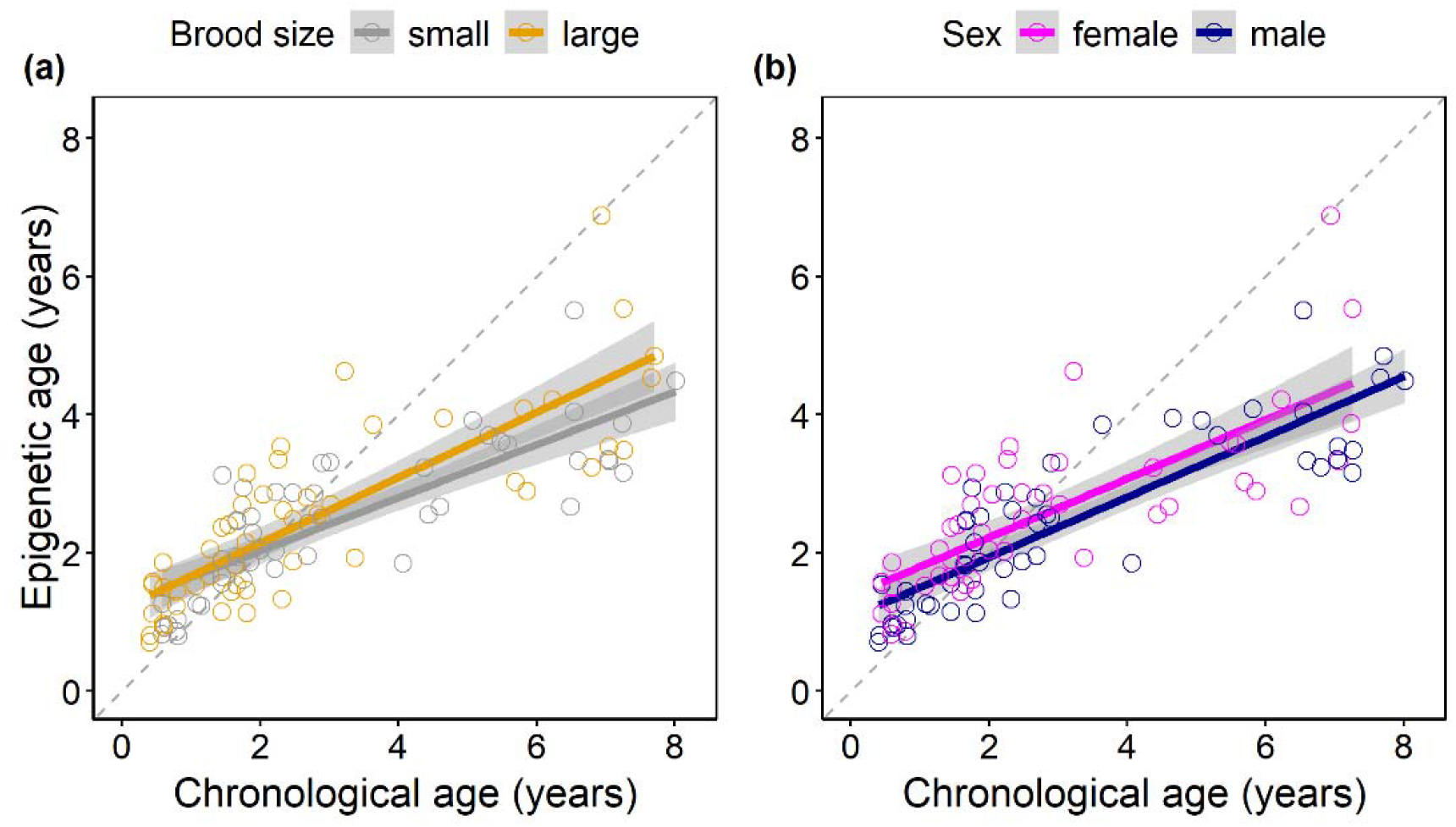
Relationship of chronological age with epigenetic age, predicted using a leave-one-individual-out cross-validation (LOIOCV) approach on 100 zebra finch samples and trained on logarithmic transformation of age for (a) males and females and (b) birds raised in small and large broods. The dashed line represents the regression line if chronological and predicted age were identical (Y=X). Solid line represents the fitted linear regression and the shaded area is the 95% CI.

**Figure S8.**
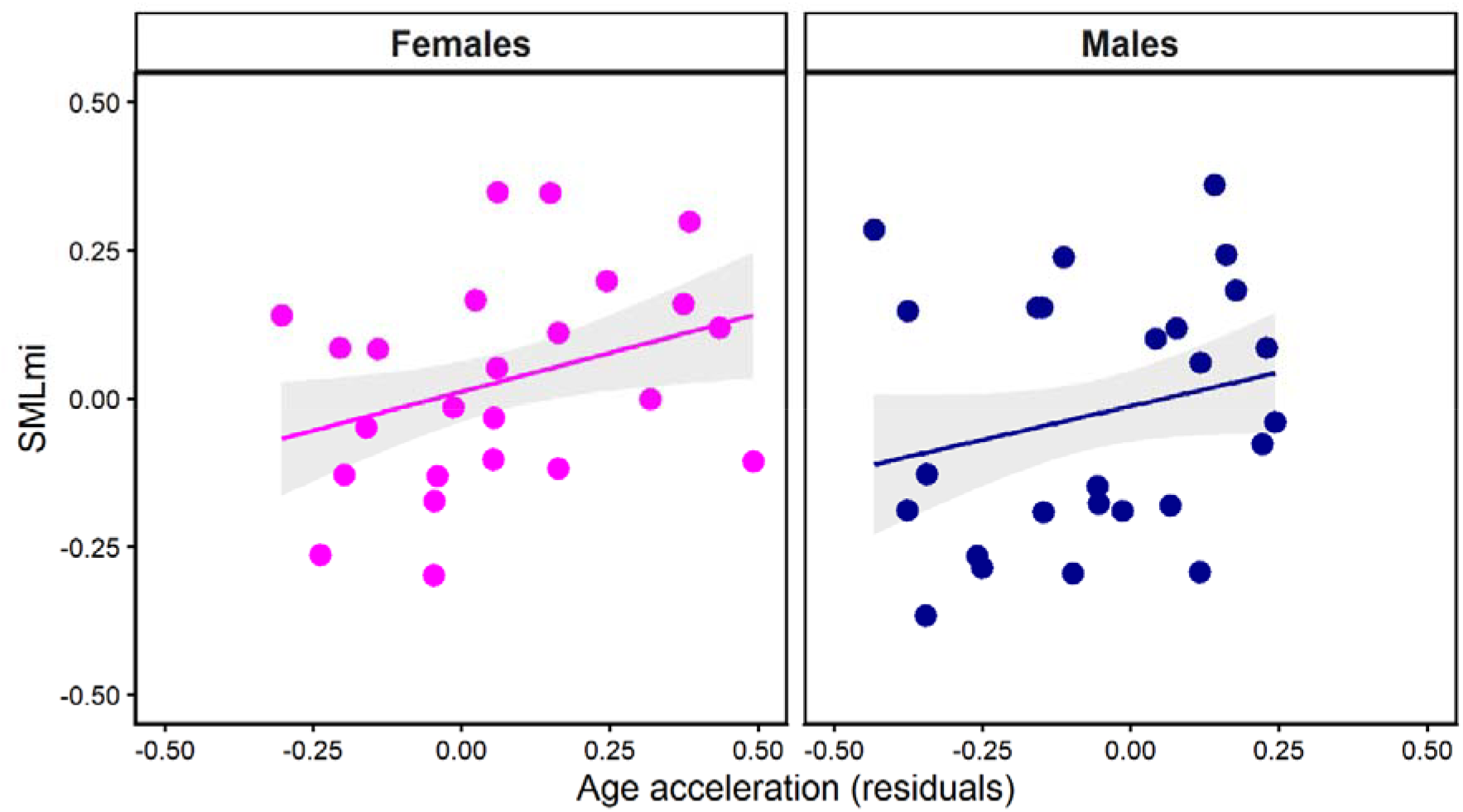
Correlation between the average per individual of the previously developed Supervised Machine Learning methylation index (SMLmi) and average epigenetic age acceleration per individual as measured by the epigenetic clock trained on the logarithmic transformation of age (N=50) separately for females and males. The solid lines represent the fitted linear regressions and the shaded areas are the 95% CI.

**Table S1.** Chromosome, position and Pearson’s correlation coefficient between chronological age and DNA methylation for the CpG sites in the EWAS of chronological age, epigenetic clock trained on the logarithmic transformation of age, epigenetic clock of age only for females and only for males.

| Chromosome | Position | Correlation<br>DNA<br>methylation<br>and age | EWAS | Clock<br>log age | Clock<br>females | Clock<br>males |
| --- | --- | --- | --- | --- | --- | --- |
| 1 | 1756530 | -0.385 |  |  |  | □ |
| 1 | 19509811 | -0.371 |  | □ | □ |  |
| 1 | 19945245 | 0.242 |  |  |  | □ |
| 1 | 20011505 | 0.254 |  |  |  | □ |
| 1 | 20874471 | -0.332 |  |  |  | □ |
| 1 | 27253521 | -0.23 |  |  | □ |  |
| 1 | 30525365 | -0.338 |  |  |  | □ |
| 1 | 33395257 | -0.474 | □ |  |  |  |
| 1 | 38973754 | -0.284 |  | □ |  |  |
| 1 | 40047616 | -0.318 |  |  |  | □ |
| 1 | 40520724 | -0.422 |  | □ |  |  |
| 1 | 41758324 | 0.138 |  |  |  | □ |
| 1 | 44676479 | -0.339 |  | □ |  |  |
| 1 | 47326980 | -0.236 |  |  | □ |  |
| 1 | 48614966 | -0.319 |  | □ |  |  |
| 1 | 49208569 | -0.38 |  | □ |  |  |
| 1 | 50827488 | -0.429 |  | □ |  |  |
| 1 | 59011852 | -0.331 |  |  |  | □ |
| 1 | 61209740 | 0.227 |  |  |  | □ |
| 1 | 67369179 | -0.411 |  | □ |  | □ |
| 1 | 74176432 | -0.47 | □ | □ | □ |  |
| 1 | 74819010 | -0.418 |  |  |  | □ |
| 1 | 76762928 | -0.237 |  |  |  | □ |
| 1 | 78312123 | -0.213 |  |  |  | □ |
| 1 | 80182749 | -0.388 |  | □ |  |  |
| 1 | 85947234 | -0.291 |  | □ |  |  |
| 1 | 89367296 | -0.288 |  |  |  | □ |
| 1 | 91995868 | -0.242 |  |  | □ |  |
| 1 | 93740767 | -0.285 |  |  |  | □ |
| 1A | 14723312 | -0.341 |  | □ |  |  |
| 1A | 18837571 | -0.318 |  |  | □ |  |
| 1A | 22691933 | -0.347 |  |  |  | □ |
| 1A | 25027881 | -0.168 |  |  |  | □ |
| 1A | 25252895 | -0.338 |  |  | □ |  |
| 1A | 25652399 | -0.297 |  |  |  | □ |
| 1A | 28098400 | -0.309 |  | □ |  |  |
| 1A | 32554112 | 0.246 |  |  |  | □ |
| 1A | 35567382 | -0.271 |  |  | □ |  |
| 1A | 38433842 | -0.257 |  |  |  | □ |
| 1A | 41902476 | -0.321 |  |  |  | □ |
| 1A | 42834540 | -0.298 |  |  |  | □ |
| 1A | 42913568 | -0.143 |  |  |  | □ |
| 1A | 43268990 | -0.503 | □ | □ |  | □ |
| 1A | 43274877 | -0.567 | □ | □ | □ | □ |
| 1A | 43274925 | -0.492 | □ | □ | □ | □ |
| 1A | 43274936 | -0.448 |  | □ |  | □ |
| 1A | 43275013 | -0.485 | □ | □ |  | □ |
| 1A | 43286149 | -0.428 |  | □ |  |  |
| 1A | 45421571 | -0.423 |  | □ |  |  |
| 1A | 45494604 | -0.389 |  | □ |  |  |
| 1A | 47159556 | -0.293 |  | □ |  |  |
| 1A | 56200897 | 0.367 |  |  |  | □ |
| 1A | 57860641 | 0.408 |  | □ |  |  |
| 1A | 58835186 | -0.333 |  | □ |  |  |
| 1A | 58960358 | -0.489 | □ | □ |  |  |
| 1A | 59884003 | 0.295 |  | □ |  |  |
| 1A | 60042208 | -0.38 |  | □ |  |  |
| 1A | 60042209 | -0.44 |  | □ |  |  |
| 1A | 60043522 | -0.471 | □ | □ |  | □ |
| 1A | 60043523 | -0.472 | □ | □ |  | □ |
| 1A | 60050548 | -0.477 | □ |  |  |  |
| 1A | 60062876 | -0.332 |  |  |  | □ |
| 1A | 60064464 | -0.54 | □ | □ |  | □ |
| 1A | 62888768 | -0.37 |  | □ |  |  |
| 1A | 62955904 | -0.293 |  |  | □ |  |
| 1A | 65221816 | -0.432 |  | □ |  |  |
| 1A | 66255373 | 0.251 |  | □ |  |  |
| 1A | 7714619 | -0.434 |  | □ |  |  |
| 2 | 1.06E+08 | 0.289 |  |  |  | □ |
| 2 | 1.06E+08 | -0.041 |  |  | □ |  |
| 2 | 1.08E+08 | 0.501 | □ | □ |  | □ |
| 2 | 1.08E+08 | -0.308 |  | □ |  |  |
| 2 | 1.15E+08 | 0.314 |  | □ |  |  |
| 2 | 1.18E+08 | -0.293 |  | □ |  |  |
| 2 | 1.19E+08 | 0.378 |  |  |  | □ |
| 2 | 1.21E+08 | 0.288 |  | □ |  |  |
| 2 | 1.24E+08 | 0.253 |  | □ |  |  |
| 2 | 1.29E+08 | -0.36 |  |  |  | □ |
| 2 | 1.34E+08 | -0.457 |  | □ |  |  |
| 2 | 1.35E+08 | -0.367 |  | □ |  |  |
| 2 | 1.4E+08 | -0.257 |  |  | □ |  |
| 2 | 1.45E+08 | 0.246 |  | □ |  |  |
| 2 | 1.52E+08 | 0.275 |  |  |  | □ |
| 2 | 15247453 | -0.311 |  | □ |  |  |
| 2 | 18115950 | 0.377 |  |  |  | □ |
| 2 | 18407925 | -0.459 |  |  |  | □ |
| 2 | 27143862 | -0.271 |  |  |  | □ |
| 2 | 30482632 | 0.502 | □ | □ | □ | □ |
| 2 | 32112422 | -0.328 |  | □ |  |  |
| 2 | 33615309 | -0.324 |  |  | □ |  |
| 2 | 36234169 | -0.342 |  | □ |  |  |
| 2 | 36352402 | 0.308 |  |  |  | □ |
| 2 | 36960156 | -0.413 |  | □ |  |  |
| 2 | 37301113 | -0.407 |  | □ |  |  |
| 2 | 41013552 | -0.348 |  | □ |  |  |
| 2 | 48398010 | -0.28 |  |  |  | □ |
| 2 | 53653059 | -0.353 |  | □ |  |  |
| 2 | 57923427 | -0.352 |  |  |  | □ |
| 2 | 59193460 | -0.352 |  |  |  | □ |
| 2 | 60190296 | -0.309 |  | □ |  |  |
| 2 | 60593395 | 0.32 |  |  |  | □ |
| 2 | 65746989 | 0.395 |  | □ |  |  |
| 2 | 71813351 | -0.34 |  |  | □ |  |
| 2 | 74249101 | 0.41 |  |  |  | □ |
| 2 | 74500507 | -0.407 |  | □ |  |  |
| 2 | 78235868 | -0.364 |  |  |  | □ |
| 2 | 79166030 | -0.386 |  |  |  | □ |
| 2 | 80453580 | -0.278 |  |  |  | □ |
| 2 | 83165815 | -0.298 |  |  |  | □ |
| 2 | 83789932 | 0.125 |  |  |  | □ |
| 2 | 84580587 | -0.357 |  |  | □ |  |
| 2 | 86475228 | -0.412 |  | □ |  |  |
| 2 | 86505822 | -0.39 |  | □ |  |  |
| 2 | 91390897 | -0.181 |  |  |  | □ |
| 2 | 95797026 | -0.363 |  |  |  | □ |
| 2 | 98654623 | 0.299 |  | □ |  |  |
| 3 | 1.11E+08 | -0.221 |  |  |  | □ |
| 3 | 1.12E+08 | 0.318 |  |  |  | □ |
| 3 | 1.12E+08 | 0.425 |  |  |  | □ |
| 3 | 1.12E+08 | 0.507 | □ |  |  | □ |
| 3 | 1.12E+08 | 0.377 |  |  |  | □ |
| 3 | 23808231 | -0.415 |  | □ |  |  |
| 3 | 23868095 | -0.263 |  |  |  | □ |
| 3 | 24991787 | 0.366 |  |  |  | □ |
| 3 | 26159693 | -0.298 |  |  |  | □ |
| 3 | 31675569 | -0.314 |  |  |  | □ |
| 3 | 31935871 | -0.419 |  | □ |  |  |
| 3 | 33873337 | -0.307 |  |  |  | □ |
| 3 | 36451990 | 0.293 |  | □ |  |  |
| 3 | 39246302 | -0.303 |  |  |  | □ |
| 3 | 40318120 | 0.334 |  |  |  | □ |
| 3 | 40874668 | -0.332 |  |  |  | □ |
| 3 | 41432632 | -0.137 |  |  | □ |  |
| 3 | 42046462 | -0.363 |  | □ |  |  |
| 3 | 44611420 | -0.421 |  |  |  | □ |
| 3 | 45280842 | 0.316 |  | □ |  |  |
| 3 | 4845539 | -0.23 |  |  |  | □ |
| 3 | 53554349 | 0.284 |  |  |  | □ |
| 3 | 5421620 | -0.364 |  |  |  | □ |
| 3 | 5504530 | 0.377 |  | □ |  |  |
| 3 | 57281964 | -0.397 |  |  |  | □ |
| 3 | 57909722 | -0.314 |  |  | □ |  |
| 3 | 59826575 | -0.188 |  |  |  | □ |
| 3 | 63437713 | 0.388 |  | □ |  |  |
| 3 | 66008234 | 0.352 |  | □ |  |  |
| 3 | 66748273 | 0.239 |  | □ |  |  |
| 3 | 66977466 | -0.18 |  |  |  | □ |
| 3 | 67346615 | 0.381 |  | □ |  |  |
| 3 | 67491744 | -0.238 |  |  |  | □ |
| 3 | 69157245 | 0.366 |  | □ |  |  |
| 3 | 74109696 | 0.239 |  |  | □ |  |
| 3 | 74591447 | -0.354 |  | □ |  |  |
| 3 | 75640604 | -0.298 |  | □ |  |  |
| 3 | 75752533 | -0.289 |  | □ |  |  |
| 3 | 75930026 | 0.311 |  |  |  | □ |
| 3 | 76175996 | -0.3 |  |  |  | □ |
| 3 | 77346911 | 0.188 |  |  |  | □ |
| 3 | 78669570 | 0.153 |  |  |  | □ |
| 3 | 86267646 | 0.317 |  | □ |  |  |
| 3 | 93364758 | 0.275 |  |  |  | □ |
| 3 | 99105229 | 0.322 |  |  |  | □ |
| 4 | 14390472 | -0.392 |  |  |  | □ |
| 4 | 17655114 | -0.356 |  |  |  | □ |
| 4 | 19003538 | -0.264 |  |  |  | □ |
| 4 | 24498070 | 0.288 |  |  |  | □ |
| 4 | 27589659 | -0.219 |  |  |  | □ |
| 4 | 27743212 | 0.262 |  |  |  | □ |
| 4 | 29746767 | -0.315 |  |  |  | □ |
| 4 | 29905507 | -0.325 |  |  |  | □ |
| 4 | 30061931 | 0.286 |  | □ |  |  |
| 4 | 31518779 | -0.439 |  |  |  | □ |
| 4 | 33128780 | -0.37 |  | □ |  | □ |
| 4 | 43891624 | -0.407 |  | □ |  |  |
| 4 | 45091977 | -0.35 |  | □ |  | □ |
| 4 | 45448908 | 0.478 | □ | □ |  | □ |
| 4 | 54548473 | -0.433 |  |  |  | □ |
| 4 | 56695096 | 0.271 |  |  |  | □ |
| 4 | 59139404 | -0.372 |  |  | □ |  |
| 4 | 59332149 | -0.379 |  |  |  | □ |
| 4 | 9075181 | -0.405 |  |  |  | □ |
| 4A | 16651377 | -0.481 | □ |  |  |  |
| 4A | 6644427 | -0.465 | □ | □ |  | □ |
| 4A | 6644494 | -0.451 |  | □ |  | □ |
| 4A | 9522855 | -0.291 |  |  |  | □ |
| 5 | 10300745 | 0.327 |  | □ |  |  |
| 5 | 16271364 | 0.267 |  |  |  | □ |
| 5 | 1926 | -0.287 |  | □ |  |  |
| 5 | 34358961 | -0.227 |  |  | □ |  |
| 5 | 36234459 | -0.151 |  |  |  | □ |
| 5 | 37580299 | -0.431 |  |  |  | □ |
| 5 | 47215701 | -0.49 | □ | □ |  | □ |
| 5 | 47445078 | -0.4 |  | □ |  |  |
| 5 | 48338006 | -0.265 |  |  |  | □ |
| 5 | 48467946 | -0.382 |  | □ |  |  |
| 5 | 49262191 | -0.303 |  |  |  | □ |
| 5 | 5100686 | 0.324 |  |  |  | □ |
| 5 | 946584 | -0.486 | □ | □ |  | □ |
| 5 | 9499105 | -0.362 |  | □ |  |  |
| 5 | 9509157 | 0.345 |  | □ |  |  |
| 6 | 14084750 | -0.299 |  |  |  | □ |
| 6 | 1841278 | -0.351 |  |  |  | □ |
| 6 | 28576688 | -0.213 |  |  |  | □ |
| 6 | 35575074 | 0.258 |  |  |  | □ |
| 6 | 6492037 | -0.454 |  | □ |  | □ |
| 6 | 6492038 | -0.516 | □ |  |  |  |
| 7 | 11107602 | -0.239 |  |  |  | □ |
| 7 | 13316430 | -0.277 |  |  |  | □ |
| 7 | 15381973 | -0.274 |  |  |  | □ |
| 7 | 24217000 | -0.323 |  |  |  | □ |
| 7 | 37901982 | 0.239 |  |  |  | □ |
| 8 | 14714587 | -0.413 |  |  |  | □ |
| 8 | 15387871 | -0.293 |  |  | □ |  |
| 8 | 16094979 | -0.2 |  |  | □ |  |
| 8 | 25629656 | -0.231 |  |  | □ |  |
| 8 | 2597538 | -0.263 |  |  |  | □ |
| 8 | 27251539 | -0.388 |  | □ |  |  |
| 8 | 2727206 | 0.237 |  | □ |  |  |
| 8 | 29575960 | 0.406 |  | □ |  |  |
| 8 | 6155619 | 0.369 |  | □ |  |  |
| 8 | 6156552 | 0.378 |  | □ |  |  |
| 8 | 6156976 | 0.364 |  | □ |  |  |
| 8 | 8942463 | -0.251 |  |  |  | □ |
| 9 | 19106745 | 0.162 |  |  |  | □ |
| 9 | 23084331 | -0.384 |  | □ |  |  |
| 9 | 23840109 | -0.29 |  |  |  | □ |
| 9 | 6442908 | 0.305 |  |  | □ |  |
| 9 | 7426004 | -0.337 |  |  |  | □ |
| 10 | 175141 | -0.201 |  |  |  | □ |
| 10 | 20329103 | -0.221 |  |  |  | □ |
| 10 | 20482402 | -0.383 |  |  |  | □ |
| 10 | 367623 | -0.38 |  | □ |  | □ |
| 10 | 367686 | -0.407 |  | □ |  | □ |
| 10 | 367743 | -0.417 |  | □ |  | □ |
| 10 | 383012 | -0.348 |  | □ | □ |  |
| 10 | 5345014 | -0.377 |  | □ |  |  |
| 11 | 12497135 | 0.383 |  | □ |  |  |
| 11 | 14546445 | 0.338 |  | □ |  |  |
| 11 | 17137158 | 0.297 |  | □ |  |  |
| 11 | 3382229 | -0.373 |  | □ |  |  |
| 12 | 11877344 | -0.389 |  | □ |  |  |
| 12 | 4169942 | 0.402 |  | □ |  |  |
| 12 | 5398190 | -0.353 |  |  |  | □ |
| 12 | 5928521 | -0.312 |  |  |  | □ |
| 13 | 18024066 | -0.397 |  | □ |  |  |
| 13 | 2502539 | -0.306 |  |  |  | □ |
| 14 | 1079720 | -0.287 |  |  |  | □ |
| 14 | 1286207 | 0.215 |  |  |  | □ |
| 14 | 41880 | -0.356 |  |  |  | □ |
| 15 | 13322302 | 0.359 |  | □ |  |  |
| 15 | 5196275 | -0.301 |  |  |  | □ |
| 15 | 6226991 | 0.351 |  |  |  | □ |
| 17 | 11219132 | -0.363 |  | □ |  |  |
| 17 | 11251539 | -0.258 |  |  |  | □ |
| 18 | 11037427 | -0.549 | □ | □ |  | □ |
| 18 | 11037867 | -0.5 | □ |  |  |  |
| 18 | 2635069 | 0.207 |  |  |  | □ |
| 19 | 121327 | -0.207 |  |  |  | □ |
| 19 | 212094 | 0.256 |  | □ |  |  |
| 19 | 3959895 | 0.264 |  |  |  | □ |
| 19 | 6278327 | -0.384 |  |  |  | □ |
| 20 | 14438453 | 0.303 |  | □ |  |  |
| 20 | 14676502 | -0.333 |  |  |  | □ |
| 20 | 1845 | -0.292 |  |  |  | □ |
| 21 | 7168673 | 0.397 |  |  |  | □ |
| 22 | 23290 | 0.357 |  | □ |  |  |
| 23 | 2129 | -0.293 |  |  | □ |  |
| 24 | 199828 | 0.323 |  |  |  | □ |
| 24 | 56935 | 0.253 |  |  |  | □ |
| 24 | 608433 | -0.306 |  |  |  | □ |
| 24 | 84345 | -0.307 |  | □ |  |  |
| 28 | 2238776 | -0.267 |  |  |  | □ |
| 28 | 534200 | -0.1 |  |  |  | □ |
| 28 | 5502931 | 0.406 |  | □ |  |  |
| 28 | 5814964 | -0.344 |  | □ |  |  |
| 32 | 2029323 | 0.572 | □ | □ |  | □ |
| 32 | 2029337 | 0.341 |  |  |  | □ |
| 32 | 2040318 | 0.314 |  |  |  | □ |
| 32 | 2042832 | 0.36 |  |  |  | □ |
| 35 | 1838443 | 0.249 |  |  |  | □ |
| 35 | 1840829 | -0.503 | □ | □ |  |  |
| 35 | 1840859 | -0.474 | □ |  |  |  |
| 35 | 1842241 | -0.152 |  |  | □ |  |
| 35 | 1843870 | -0.478 | □ |  |  |  |
| 35 | 1848109 | -0.47 | □ |  |  | □ |
| 35 | 1912073 | 0.267 |  |  |  | □ |
| Z | 58226856 | -0.498 | □ |  |  |  |
| Z | 6438489 | -0.47 | □ |  |  |  |
| Z | 74482930 | -0.237 |  |  |  | □ |

**Table S2.**
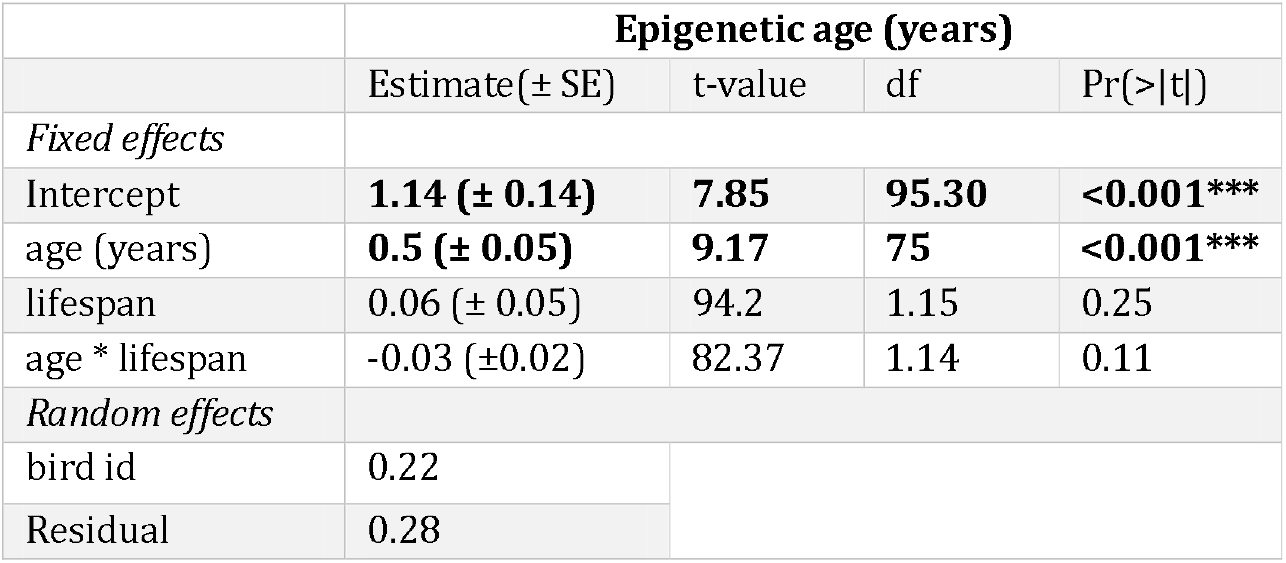
Epigenetic age in relation to age and lifespan. Epigenetic clock was trained on logarithmic transformation of age and lifespan was mean centered.

**Table S3.** Results of linear mixed effects models on the effects of delta and mean age acceleration on epigenetic age from the epigenetic clock trained on logarithmic transformation of age separately for females and males.

| **Females** |  |  |  |  |
| --- | --- | --- | --- | --- |
|  | **Epigenetic age (years)** |  |  |  |
|  | Estimate(± SE) | t-value | df | Pr(>\|t\|) |
| *Fixed effects* |  |  |  |  |
| Intercept | -0.01(± 0.04) | -0.32 | 22 | 0.76 |
| delta age acceleration | -0.04 (± 0.13) | -0.29 | 23 | 0.77 |
| mean age acceleration | 0.26 (± 0.16) | 1.63 | 22 | 0.12 |
| *Random effects* |  |  |  |  |
| bird id | 0.04 |  |  |  |
| Residual | 0.005 |  |  |  |
| **Males** |  |  |  |  |
|  | **Epigenetic age (years)** |  |  |  |
|  | Estimate(± SE) | t-value | df | Pr(>\|t\|) |
| *Fixed effects* |  |  |  |  |
| Intercept | -0.13 (± 0.04) | -0.3 | 24 | 0.77 |
| delta age acceleration | -0.009 (± 0.08) | -0.12 | 25 | 0.91 |
| mean age acceleration | 0.23 (± 0.19) | 1.41 | 24 | 0.27 |
| *Random effects* |  |  |  |  |
| bird id | 0.16 |  |  |  |
| Residual | 0.30 |  |  |  |

**Table S4.** Results of linear mixed effects model on the effects of brood size and sex on epigenetic age from the epigenetic clock trained on logarithmic transformation of age. Reference categories are small broods and females.

|  | **Epigenetic age (years)** |  |  |  |
| --- | --- | --- | --- | --- |
|  | Estimate (± SE) | t-value | df | Pr(>\|t\|) |
| *Fixed effects* |  |  |  |  |
| Intercept | **1.19 (± 0.21)** | **75.15** | **5.69** | **<0.001\*\*\*** |
| age (years) | **0.4 (± 0.04)** | **71.32** | **10.38** | **<0.001\*\*\*** |
| brood size | 0.3 (± 0.28) | 71.38 | 1.08 | 0.283 |
| sex | 0.03 (±0.23) | 44.07 | 0.13 | 0.895 |
| brood size * sex | -0.6 (±0.32) | 44.37 | -1.89 | 0.066 |
| age * brood size | 0.06 (±0.06) | 78.24 | 1.14 | 0.260 |
| *Random effects* |  |  |  |  |
| bird id | 0.16 |  |  |  |
| Residual | 0.30 |  |  |  |

## References

Akalin, A., Kormaksson, M., Li, S., Garrett-Bakelman, F. E., Figueroa, M. E., Melnick, A., & Mason, C. E. (2012). MethylKit: a comprehensive R package for the analysis of genome-wide DNA methylation profiles. Genome Biology, 13(10), 1–9. 10.1186/gb-2012-13-10-R87

Andrews, S. (2010). FastQC: a quality control tool for high throughput sequence data.

Aviv, A., & Verhulst, S. (2025). Telomeres in Space. Aging Cell, 24(3), 1–4. 10.1111/acel.70030

Bates, D., Mächler, M., Bolker, B. M., & Walker, S. C. (2015). Fitting linear mixed-effects models using lme4. Journal of Statistical Software, 67(1), v1.1-21. 10.18637/jss.v067.i01

Benjamini, Y., & Hochberg, Y. (1995). Controlling the false discovery ratelz: A practical and powerful approach to multiple testing. Journal of the Royal Statistical Society, 57(1), 289–300.

Berdyshev, G. D., Korotaev, G. K., Boiarskikh, G. V, & Vaniushin, B. F. (1967). Nucleotide composition of DNA and RNA from somatic tissues of humpback and its changes during spawning. Biokhimiia (Moscow, Russia), 32(5), 988—993. http://europepmc.org/abstract/MED/5628601

Berman, B. P., Weisenberger, D. J., Aman, J. F., Hinoue, T., Ramjan, Z., Liu, Y., Noushmehr, H., Lange, C. P. E., VanDijk, C. M., Tollnaar, R. A. E. M., VanDenBerg, D., & Laird, P. W. (2015). Regions of focal DNA hypermethylation and long-range hypomethylation in colorectal cancer coincide with nuclear lamina–associated domains. Nature Genetics, 44(1), 40–46. 10.1016/j.coph.2007.10.002.Taste

Bland, J. M., & Altman, D. G. (1994). Statistic Notes: Regression towards the mean. BMJ, 308, 1499.

Bogdanović, O., & Lister, R. (2017). DNA methylation and the preservation of cell identity. Current Opinion in Genetics and Development, 46, 9–14. 10.1016/j.gde.2017.06.007

Bollati, V., Schwartz, J., Wright, R., Litonjua, A., Tarantini, L., Suh, H., Sparrow, D., Vokonas, P., & Baccarelli, A. (2009). Decline in genomic DNA methylation through aging in a cohort of elderly subjects. Mechanisms of Ageing and Development, 130(4), 234–239. 10.1016/j.mad.2008.12.003

Briga, M., Koetsier, E., Boonekamp, J. J., Jimeno, B., & Verhulst, S. (2017a). Food availability affects adult survival trajectories depending on early developmental conditions. Proceedings of the Royal Society B: Biological Sciences, 284(1846), 20162287. 10.1098/rspb.2016.2287

Briga, M., Koetsier, E., Boonekamp, J. J., Jimeno, B., & Verhulst, S. (2017b). Food availability affects adult survival trajectories depending on early developmental conditions. Proceedings of the Royal Society B: Biological Sciences, 284(1846), 20162287. 10.1098/rspb.2016.2287

Casellas, J., & Medrano, J. F. (2008). Lack of Socs2 expression reduces lifespan in high-growth mice. Age, 30(4), 245–249. 10.1007/s11357-008-9064-1

De Paoli-Iseppi, R., Deagle, B. E., Polanowski, A. M., McMahon, C. R., Dickinson, J. L., Hindell, M. A., & Jarman, S. N. (2019). Age estimation in a long-lived seabird (Ardenna tenuirostris) using DNA methylation-based biomarkers. Molecular Ecology Resources, 19(2), 411–425. 10.1111/1755-0998.12981

Dobson, A. J., Boulton-McDonald, R., Houchou, L., Svermova, T., Ren, Z., Subrini, J., Vazquez-Prada, M., Hoti, M., Rodriguez-Lopez, M., Ibrahim, R., Gregoriou, A., Gkantiragas, A., Bähler, J., Ezcurra, M., & Alic, N. (2019). Longevity is determined by ETS transcription factors in multiple tissues and diverse species. PLoS Genetics, 15(7). 10.1371/journal.pgen.1008212

Drewelies, J., Homann, J., Vetter, V. M., Düzel, S., Kühn, S., Deecke, L., Steinhagen-Thiessen, E., Jawinski, P., Markett, S., Lindenberger, U., Lill, C. M., Bertram, L., Demuth, I., & Gerstorf, D. (2025). There Are Multiple Clocks That Time Us: Cross-Sectional and Longitudinal Associations Among 14 Alternative Indicators of Age and Aging. The Journals of Gerontology-Series A: Biological Sciences and Medical Sciences, 80(6). 10.1093/gerona/glae244

Elster, D., & von Eyss, B. (2020). Hippo signaling in regeneration and aging. Mechanisms of Ageing and Development, 189. 10.1016/j.mad.2020.111280

Ewels, P., Magnusson, M., Lundin, S., & Käller, M. (2016). MultiQC: Summarize analysis results for multiple tools and samples in a single report. Bioinformatics, 32(19), 3047–3048. 10.1093/bioinformatics/btw354

Fernandes Martins, T. L. (2004). Sex-specific growth rates in zebra finch nestlings: A possible mechanism for sex ratio adjustment. Behavioral Ecology, 15(1), 174–180. 10.1093/beheco/arg094

Friedman, J., Hastie, T., & Tibshirani, R. (2010). Regularization Paths for Generalized Linear Models via Coordinate Descent. Journal of Statistical Software, 33(1), 1–22. 10.18637/jss.v033.i01

Gerber, L., Peters, K. J., King, S. L., Allen, S. J., Connor, R. C., Forbes, O., Holmes, K. G., Kearns, A. M., Willems, E. P., Krützen, M., & Rollins, L. A. (2025). Social bonds decrease epigenetic age in male bottlenose dolphins. Communications Biology, 81(1765), 1–8. 10.1038/s42003-025-09227-w

Gerber, L., Schrey, A. W., Anderson, S. C., Jain, E., & Liebl, A. L. (2025). Sequencing method matters: differential performance of DNA methylation data acquisition in epigenetic clock calibration. Journal of Avian Biology, 2025(5). 10.1002/jav.03498

Griffith, S. C., Crino, O. L., Andrew, S. C., Nomano, F. Y., Adkins-Regan, E., Alonso-Alvarez, C., Bailey, I. E., Bittner, S. S., Bolton, P. E., Boner, W., Boogert, N. J., Boucaud, I. C. A., Briga, M., Buchanan, K. L., Caspers, B. A., Cichoń, M., Clayton, D. F., Derégnaucourt, S., Forstmeier, W., … Williams, T. D. (2017). Variation in Reproductive Success Across Captive Populations: Methodological Differences, Potential Biases and Opportunities. Ethology, 123(1), 1–29. 10.1111/eth.12576

Gryzinska, M., Blaszczak, E., Strachecka, A., & Jezewska-Witkowska, G. (2013). Analysis of age-related global DNA methylation in chicken. Biochemical Genetics, 51(7–8), 554–563. 10.1007/s10528-013-9586-9

Haller, A., Risse, J., Sepers, B., & van Oers, K. (2025). Independent Avian Epigenetic Clocks for Ageing and Development. Molecular Ecology Resources, 25(7), 1–11. 10.1111/1755-0998.14128

Higgins-Chen, A. T., Thrush, K. L., Wang, Y., Minteer, C. J., Kuo, P. L., Wang, M., Niimi, P., Sturm, G., Lin, J., Moore, A. Z., Bandinelli, S., Vinkers, C. H., Vermetten, E., Rutten, B. P. F., Geuze, E., Okhuijsen-Pfeifer, C., van der Horst, M. Z., Schreiter, S., Gutwinski, S., … Levine, M. E. (2022). A computational solution for bolstering reliability of epigenetic clocks: implications for clinical trials and longitudinal tracking. Nature Aging, 2(7), 644–661. 10.1038/s43587-022-00248-2

Horvath, S. (2013). DNA methylation age of human tissues and cell types. Genome Biology, 16(1), 1–19. 10.1186/s13059-015-0649-6

Horvath, S., Erhart, W., Brosch, M., Ammerpohl, O., Von Schönfels, W., Ahrens, M., Heits, N., Bell, J. T., Tsai, P. C., Spector, T. D., Deloukas, P., Siebert, R., Sipos, B., Becker, T., Röcken, C., Schafmayer, C., & Hampe, J. (2014). Obesity accelerates epigenetic aging of human liver. Proceedings of the National Academy of Sciences of the United States of America, 111(43), 15538–15543. 10.1073/pnas.1412759111

Horvath, S., Lu, A. T., Haghani, A., Zoller, J. A., Li, C. Z., Lim, A. R., Brooke, R. T., Raj, K., Serres-Armero, A., Dreger, D. L., Hogan, A. N., Plassais, J., & Ostrander, E. A. (2022). DNA methylation clocks for dogs and humans. Proceedings of the National Academy of Sciences of the United States of America, 119(21). 10.1073/pnas.2120887119

Horvath, S., & Raj, K. (2018). DNA methylation-based biomarkers and the epigenetic clock theory of ageing. In Nature Reviews Genetics (Vol. 19, Number 6, pp. 371–384). Springer US. 10.1038/s41576-018-0004-3

Karpf, A. R., & Matsui, S. I. (2005). Genetic disruption of cytosine DNA methyltransferase enzymes induces chromosomal instability in human cancer cells. Cancer Research, 65(19), 8635–8639. 10.1158/0008-5472.CAN-05-1961

Koetsier, E., & Verhulst, S. (2011). A simple technique to manipulate foraging costs in seed-eating birds. The Journal of Experimental Biology, 214(Pt 8), 1225–1229. 10.1242/jeb.050336

Krueger, F., & Andrews, S. R. (2011). Bismark: A flexible aligner and methylation caller for Bisulfite-Seq applications. Bioinformatics, 27(11), 1571–1572. 10.1093/bioinformatics/btr167

Krueger, F., James, F., Ewels, P., Afyounian, E., Weinstein, M., Schuster-Boeckler, B., Hulselmans, G., & sclamons. (2023). FelixKrueger/TrimGalore: v0.6.10. Zenodo. 10.5281/zenodo.7598955

Kuo, P.-L., Moore, A. Z., Tanaka, T., Belsky, D. W., Lu, A. T.-H., Horvath, S., Bandinelli, S., & Ferrucci, L. (2026). Longitudinal changes in epigenetic clocks predict survival in the InCHIANTI cohort. Nature Aging, 6(3), 534–540. 10.1038/s43587-026-01066-6

Kwon, A., Chae, H. W., Lee, W. J., Kim, J. H., Kim, Y. J., Ahn, J., Oh, Y., & Kim, H. S. (2023). Insulin-like growth factor binding protein-3 induces senescence by inhibiting telomerase activity in MCF-7 breast cancer cells. Scientific Reports, 13(1). 10.1038/s41598-023-35291-5

Langfelder, P., & Horvath, S. (2008). WGCNA: An R package for weighted correlation network analysis. BMC Bioinformatics, 9. 10.1186/1471-2105-9-559

Langmead, B., & Salzberg, S. L. (2012). Fast gapped-read alignment with Bowtie 2. Nature Methods, 9(4), 357–359. 10.1038/nmeth.1923

Lee, M. G. (2026). The Age Illusion — Limitations of Chronologic Age in Medicine. New England Journal of Medicine, 394(13), 1249–1251. 10.1056/NEJMp2415488

Lemaître, J. F., Moorad, J., Gaillard, J. M., Maklakov, A. A., & Nussey, D. H. (2024). A unified framework for evolutionary genetic and physiological theories of aging. PLoS Biology, 22(2 February). 10.1371/journal.pbio.3002513

Letellier, E., & Haan, S. (2016). SOCS2: physiological and pathological functions. Frontiers in Bioscience, (8), 189–204. 10.2741/E760

Levine, M. E., Lu, A. T., Quach, A., Chen, B. H., Assimes, T. L., Hou, L., Baccarelli, A. A., Stewart, J. D., Li, Y., Whitsel, E. A., Wilson, G., Reiner, A. P., Aviv, A., Lohman, K., Liu, Y., & Ferrucci, L. (2018). An epigenetic biomarker of aging for lifespan and healthspan. Aging, 10(4), 573–591.

Li, A., Mueller, A., English, B., Arena, A., Vera, D., Kane, A. E., & Sinclair, D. A. (2022). Novel feature selection methods for construction of accurate epigenetic clocks. PLoS Computational Biology, 18(8). 10.1371/journal.pcbi.1009938

Li, C. Z., Haghani, A., Yan, Q., Lu, A. T., Zhang, J., Fei, Z., Ernst, J., William Yang, X., Gladyshev, V. N., Robeck, T. R., Chavez, A. S., Cook, J. A., Dunnum, J. L., Raj, K., Seluanov, A., Gorbunova, V., & Horvath, S. (2024). O R G A N I S M A L B I O L O G Y Epigenetic predictors of species maximum life span and other life-history traits in mammals. Science Advances, 10, 7273. 10.5281/ze

López-Otín, C., Blasco, M. A., Partridge, L., Serrano, M., & Kroemer, G. (2023). Hallmarks of aging: An expanding universe. Cell, 186(2), 243–278. 10.1016/j.cell.2022.11.001

Lowsky, D. J., Olshansky, S. J., Bhattacharya, J., & Goldman, D. P. (2014). Heterogeneity in healthy aging. Journals of Gerontology - Series A Biological Sciences and Medical Sciences, 69(6), 640–649. 10.1093/gerona/glt162

Lu, A. T., Quach, A., Wilson, J. G., Reiner, A. P., Aviv, A., Raj, K., Hou, L., Baccarelli, A. A., Li, Y., Stewart, J. D., Whitsel, E. A., & Themistocles, L. (2019). DNA methylation GrimAge strongly predicts lifespan and healthspan. Aging, 11(2), 303–327.

Meyer, B. S., Moiron, M., Caswara, C., Chow, W., Fedrigo, O., Formenti, G., Haase, B., Howe, K., Mountcastle, J., Uliano-Silva, M., Wood, J., Jarvis, E. D., Liedvogel, M., & Bouwhuis, S. (2023). Sex-specific changes in autosomal methylation rate in ageing common terns. Frontiers in Ecology and Evolution, 11(January), 1–9. 10.3389/fevo.2023.982443

Moore, L. D., Le, T., & Fan, G. (2013). DNA methylation and its basic function. Neuropsychopharmacology, 38(1), 23–38. 10.1038/npp.2012.112

Parrott, B. B., & Bertucci, E. M. (2019). Epigenetic Aging Clocks in Ecology and Evolution. Trends in Ecology and Evolution, 34(9), 767–770. 10.1016/j.tree.2019.06.008

R Core Team. (2023). R: A language and environment for statistical computing. (RStudio v. 4.3.2). https://www.r-project.org/

Rhie, A., McCarthy, S. A., Fedrigo, O., Damas, J., Formenti, G., Koren, S., Uliano-Silva, M., Chow, W., Fungtammasan, A., Kim, J., Lee, C., Ko, B. J., Chaisson, M., Gedman, G. L., Cantin, L. J., Thibaud-Nissen, F., Haggerty, L., Bista, I., Smith, M., … Jarvis, E. D. (2021). Towards complete and error-free genome assemblies of all vertebrate species. Nature, 592(7856), 737–746. 10.1038/s41586-021-03451-0

Rutledge, J., Oh, H., & Wyss-Coray, T. (2022). Measuring biological age using omics data. Nature Reviews Genetics. 10.1038/s41576-022-00511-7

Sheaffer, K. L., Elliott, E. N., & Kaestner, K. H. (2016). DNA hypomethylation contributes to genomic instability and intestinal cancer initiation. Cancer Prevention Research, 9(7), 534–546. 10.1158/1940-6207.CAPR-15-0349.DNA

Sheldon, E. L., Schrey, A. W., Ragsdale, A. K., & Griffith, S. C. (2018). Brood size influences patterns of DNA methylation in wild Zebra Finches (Taeniopygia guttata). Auk, 135(4), 1113–1122. 10.1642/AUK-18-61.1

Simpson, D. J., & Chandra, T. (2021). Epigenetic age prediction. In Aging Cell (Vol. 20, Number 9). John Wiley and Sons Inc. 10.1111/acel.13452

Stoffel, M. A., Nakagawa, S., & Schielzeth, H. (2017). rptR: repeatability estimation and variance decomposition by generalized linear mixed-effects models. Methods in Ecology and Evolution, 8(11), 1639–1644. 10.1111/2041-210X.12797

Tangili, M., Briga, M., & Verhulst, S. (2026). Begging efficiency rather than food received causes brood size effect on growth in zebra finches. Behaviour, 1–22. 10.1163/1568539X-bja10358

Tangili, M., Mulder, E., Jimeno, B., Briga, M., & Verhulst, S. (2026). Telomere dynamics, not absolute telomere length, predicts lifespan in adult zebra finches. Journal of Evolutionary Biology. 10.1093/jeb/voag034

Tangili, M., Palsbøll, P. J., & Verhulst, S. (2026). Inferences from epigenetic information in an ecological context: a case study of early-life environmental effects on DNA methylation in zebra finches. BioRxiv. 10.1101/2025.07.30.667588

Tangili, M., Slettenhaar, A. J., Sudyka, J., Dugdale, H. L., Pen, I., Palsbøll, P. J., & Verhulst, S. (2023a). DNA methylation markers of age(ing) in non-model animals. Molecular Ecology, 32(17), 4725–4741. 10.1111/mec.17065

Tangili, M., Slettenhaar, A. J., Sudyka, J., Dugdale, H. L., Pen, I., Palsbøll, P. J., & Verhulst, S. (2023b). DNA methylation markers of age(ing) in non-model animals. Molecular Ecology, 32(17), 4725–4741. 10.1111/mec.17065

Tangili, M., Sudyka, J., Furni, F., Palsbøll, P. J., & Verhulst, S. (2024). Dataset-Identification of age-related CpG sites from longitudinal avian methylomes. Dryad. 10.5061/dryad.wm37pvmw8

Tangili, M., Sudyka, J., Furni, F., Palsbøll, P. J., & Verhulst, S. (2025). Sex-Chromosome-Dependent Ageing in Female Heterogametic Methylomes. Molecular Ecology, 34(e70147), 1–11. 10.1111/mec.70147

Thompson, M. J., Chwia, K., Rubbi, L., Lusis, A. J., Richard, C., Srivastava, A., Korstanje, R., Churchill, G. A., Horvath, S., & Pellegrini, M. (2018). A multi - tissue full lifespan epigenetic clock for mice. Aging, 10(10), 2832–2854.

Timp, W., Bravo, H. C., McDonald, O. G., Goggins, M., Umbricht, C., Zeiger, M., Feinberg, A. P., & Irizarry, R. A. (2014). Large hypomethylated blocks as a universal defining epigenetic alteration in human solid tumors. Genome Medicine, 6(8), 1–11. 10.1186/s13073-014-0061-y

Tinbergen, J. M., & Boerlijst, M. C. (1990). Nestling Weight and Survival in Individual Great Tits (Parus major). The Journal of Animal Ecology, 59(3), 1113. 10.2307/5035

Unnikrishnan, A., Hadad, N., Masser, D. R., Jackson, J., Freeman, W. M., & Richardson, A. (2018). Revisiting the genomic hypomethylation hypothesis of aging. Annals of the New York Academy of Sciences, 1418(1), 69–79. 10.1111/nyas.13533

Vanyushin, B. F., Nemirovsky, L. E., Klimenko, V. V, Vasiliev, V. K., & Belozersky, A. N. (1973). The 5-methylcytosine in DNA of rats. Tissue and age specificity and the changes induced by hydrocortisone and other agents. Gerontologia, 19(3), 138–152.

Wilson, V. L., Smith, R. A., Ma, S., & Cutler, R. G. (1987). Genomic 5-methyldeoxycytidine decreases with age. Journal of Biological Chemistry, 262(21), 9948–9951. 10.1016/s0021-9258(18)61057-9

Wolf, S. E., & Tangili, M. (2026). Avian epigenetic clocks: state of the art and call to action. EcoEvoRxiv. 10.32942/X2239P

Xiao, F. H., Kong, Q. P., Perry, B., & He, Y. H. (2016). Progress on the role of DNA methylation in aging and longevity. Briefings in Functional Genomics, 15(6), 454–459. 10.1093/bfgp/elw009

Yalaev, B., Tyurin, A., Akhiiarova, K., & Khusainova, R. (2024). Hypomethylation of the RUNX2 Gene Is a New Potential Biomarker of Primary Osteoporosis in Men and Women. International Journal of Molecular Sciences, 25(13). 10.3390/ijms25137312

Zann, R. A. (1996). The zebra finch. A synthesis of field and laboratory studies. Oxford University Press.

Ziller, M. J., Hansen, K. D., Meissner, A., Martin, J., Biology, R., Unit, P., Hospital, M. G., & Hospital, M. G. (2015). Coverage recommendations for methylation analysis by whole genome bisulfite sequencing. Nature Methods, 12(3), 230–232. 10.1038/nmeth.3152.Coverage

